# Neural Fragility as an EEG Marker of the Seizure Onset Zone

**DOI:** 10.1101/862797

**Authors:** Adam Li, Chester Huynh, Zachary Fitzgerald, Iahn Cajigas, Damian Brusko, Jonathan Jagid, Angel Claudio, Andres Kanner, Jennifer Hopp, Stephanie Chen, Jennifer Haagensen, Emily Johnson, William Anderson, Nathan Crone, Sara Inati, Kareem Zaghloul, Juan Bulacio, Jorge Gonzalez-Martinez, Sridevi V. Sarma

## Abstract

Over 15 million epilepsy patients worldwide do not respond to drugs. Successful surgical treatment requires complete removal, or disconnection of the seizure onset zone (SOZ), brain region(s) where seizures originate. Unfortunately, surgical success rates vary between 30%-70% because no clinically validated biological marker of the SOZ exists. We develop and retrospectively validate a new EEG marker - *neural fragility* - in a retrospective analysis of 91 patients by using neural fragility of the annotated SOZ as a metric to predict surgical outcomes. Fragility predicts 43/47 surgical failures with an overall prediction accuracy of 76%, compared to the accuracy of clinicians being 48% (successful outcomes). In failed outcomes, we identify fragile regions that were untreated. When compared to 20 EEG features proposed as SOZ markers, fragility outperformed in predictive power and interpretability suggesting neural fragility as an EEG biomarker of the *SOZ*.

## 1 Introduction

Over 15 million epilepsy patients worldwide and 1 million in the US suffer from drug-resistant epilepsy (DRE) [1, 2]. DRE is defined as continued seizures despite adequate trials of two tolerated appropriately chosen anti-epileptic drugs [3]. DRE patients have an increased risk of sudden death and are frequently hospitalized, burdened by epilepsy-related disabilities, and the cost of their care is a significant contributor to the $16 billion dollars spent annually in the US treating epilepsy patients [4]. Approximately 50% of DRE patients have focal DRE, where a specific brain region or regions, termed the epileptogenic zone (EZ), is necessary and sufficient for initiating seizures and whose removal (or disconnection) results in complete abolition of seizures [5, 6]. The EZ encompasses the clinically identified seizure onset zone (SOZ) and early propagation zone (EPZ). The brain regions associated with the *SOZ* demonstrate the earliest electrophysiological changes during a seizure event, and in general precede the clinical onset of seizures; and the EPZ regions are involved at the time of the earliest clinical (semiological) manifestations during a seizure event. Successful surgical and neuromodulatory treatments can stop seizures altogether or allow them to be controlled with medications [7], but outcomes for both treatments critically depend on accurate localization of the *SOZ*.

Localizing the *SOZ* also relies on accurate placement of electrodes such that they cover the EZ, and the ability to identify abnormalities in the intracranial EEG (iEEG) channels that may correlate to the *SOZ* with the naked eye. Unfortunately, even the most experienced clinicians are challenged because epilepsy is fundamentally a network disease, which cannot be entirely defined by the current methods of localization. Abnormal connections across several channels may constitute a more effective marker of the *SOZ* [8]. Localization thus lends itself to a data-driven network-based computational approach and several electroencephalogram (EEG) algorithms have been proposed to localize the *SOZ* from recordings. Many entail investigations of the spectral power in each iEEG channel including high frequency oscillations [9], but these approaches do not consider network properties of the brain because they treat each EEG channel independently. Others have proposed graph-based analysis of iEEG [10, 11, 12, 13, 14, 15], but these approaches fail to identify internal network properties that cause seizures to occur in the first place.

We propose a new EEG marker of the *SOZ*, neural fragility, of brain regions (conceptually described in Figure 1). To create the fragility marker, we first build a personalized dynamic model of the brain network from observed iEEG signals. The model is generative in that it can accurately reconstruct the patient’s iEEG recordings [16, 17]. We then calculate which network nodes are imbalanced, meaning small impulse perturbations on the network and thus can trigger seizures (see Figure 1). Neural fragility measures the degree to which a node is imbalanced [18]. To evaluate neural fragility as a marker of the SOZ, we conducted a retrospective study using iEEG data from 91 patients treated across 5 epilepsy centers: Johns Hopkins Hospital (JHH), National Institute of Health (NIH), Cleveland Clinic (CClinic), University of Maryland Medical Center (UMMC) and Jackson Memorial Hospital, University of Miami (UMF). In the study population, all DRE patients underwent invasive iEEG monitoring followed by surgical resection or laser ablation of the *SOZ* (44 success and 47 failure outcome). We demonstrate that neural fragility is higher/lower in electrode contacts within clinically annotated SOZs for success/failure patients. In addition, we compare fragility of iEEG nodes to 6 frequency-based and 14 graph theoretic features in a 10-fold nested-cross validation. Neural fragility has an area under the curve (AUC) discrimination score of 0.88 +/−0.064, which is 13% better compared to the next best feature. In addition, it has a high degree of interpretability, which we demonstrate by computing an interpretability ratio suggesting that the spatiotemporal heatmaps of neural fragility are a robust iEEG biomarker of the *SOZ* that can incorporate seamlessly into the clinical workflow.

**Figure 1:**
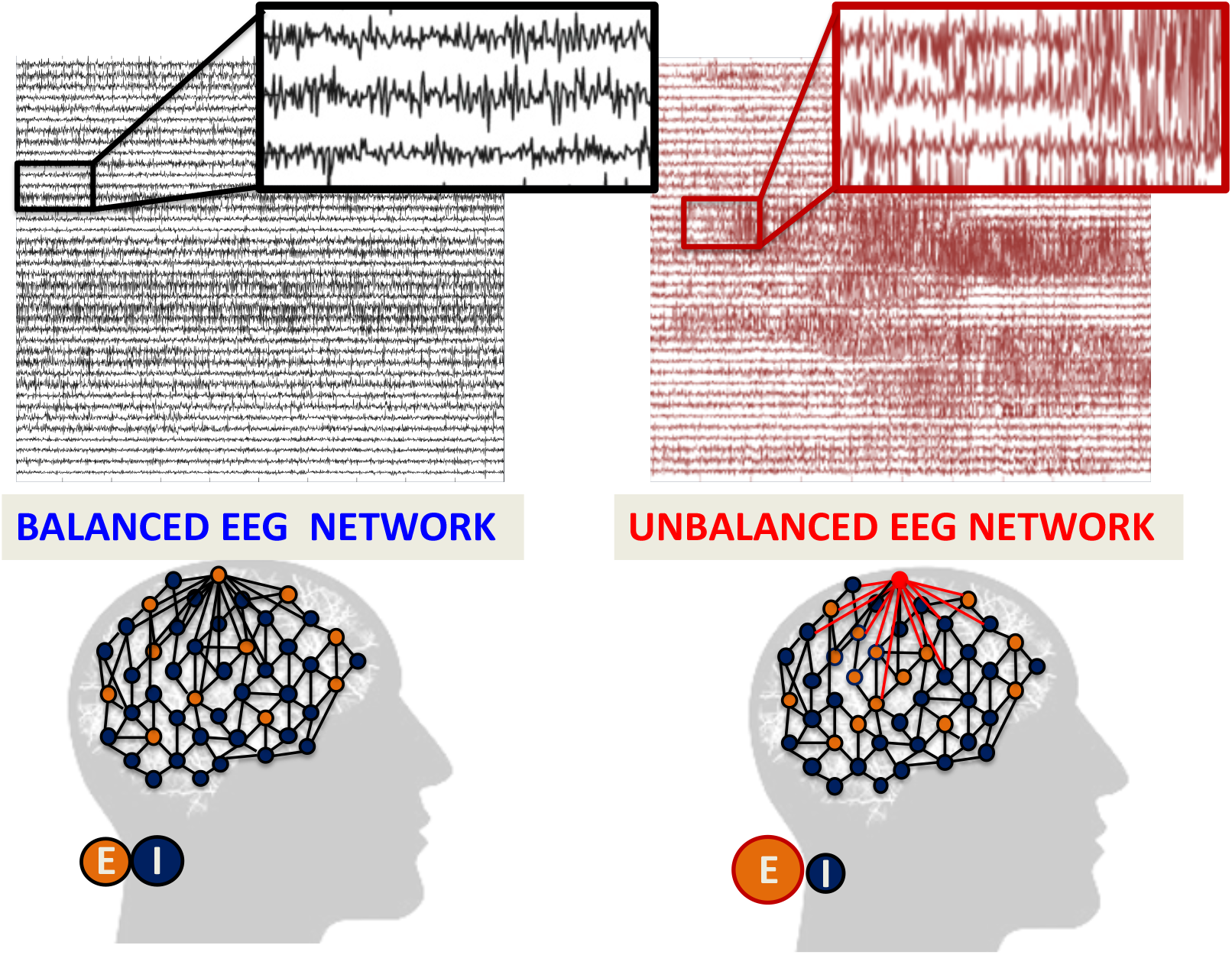
Intuition of neural fragility - unbalanced and balanced networks. **(Top)** iEEG traces in between seizures (left) and during a seizure (right). **(Bottom)** network schematic showing change in connectivity (right) in fragile node that causes seizure. Describes qualitatively the concept of neural fragility in the context of a dynamical iEEG network, with nodes representing excitatory (*E*) and inhibitory (*I*) population of neurons. From a dynamical systems point of view, such imbalance arises from a few fragile nodes causing instability of the network in the form of over-excitation, or under-inhibition. We define fragility of a network node to be the minimum-energy perturbation applied to the node’s weights on its neighbors before rendering the network unstable [18, 16]. In systems theory, stable systems return to a baseline condition when a node is perturbed. In contrast, unstable systems can oscillate and grow when a node is perturbed. In the context of epilepsy, a fragile node is one that requires a smaller perturbation to lead to seizure activity. Fragility theory can be modeled in the context of linear dynamical systems: *x*(*t* + 1) = *Ax*(*t*). Perturbing the columns of the A matrix will alter dynamical connections of a particular node (i.e. that column) on its neighbors, resulting in an imbalanced network.

## 2 Results

There is no clinically validated biomarker of the *SOZ*, yet it is an important component for the localization of the underlying EZ [19]. This presents a serious challenge for clinicians to accurately localize the *SOZ* and has led to surgical success rates to vary between 30-70% despite large brain regions being removed [20]. With no biomarker available, clinicians conduct extensive evaluations with neuroimaging, clinical testing, and visual inspection of EEG recordings (Figure 3). When non-invasive tests are inconclusive, patients undergo intracranial monitoring, during which iEEG electrodes are either placed directly on the cortex or implanted into the brain. iEEG provides high temporal resolution data for clinicians to visually detect abnormal activity, such as spikes and high frequency bursts, in between seizures (interictal) and during seizures (ictal). Specifically, clinicians attempt to identify electrodes involved in the *SOZ* and early spread [21]. Surgical resection is then performed on the basis of this hypothesis, unless the region overlaps with eloquent cortex.

**Figure 3:**
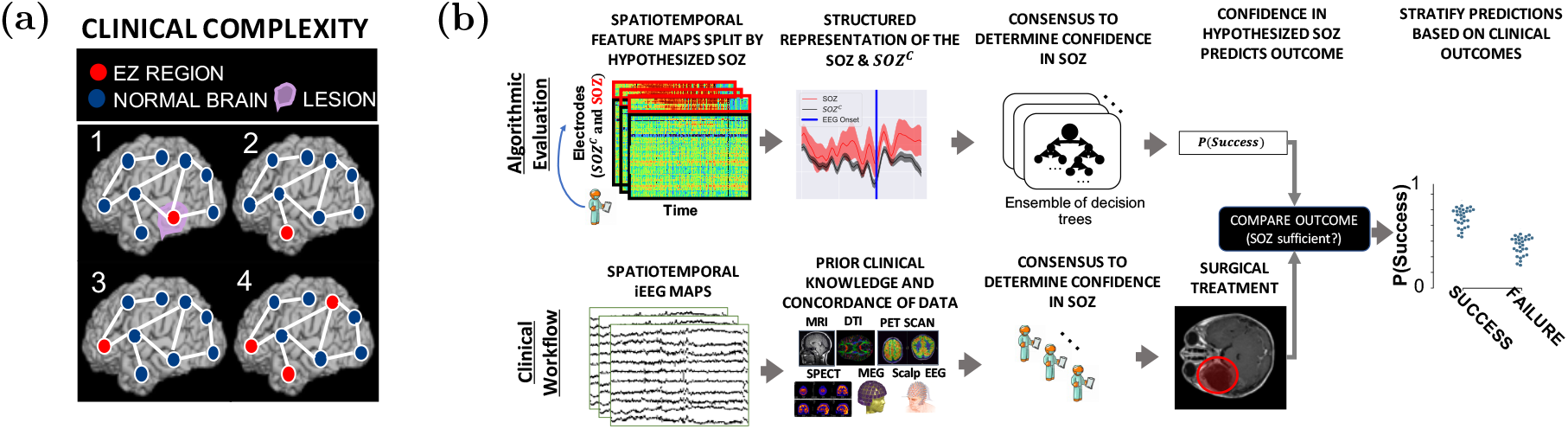
Clinical complexity and our experimental paradigm. **(a)** Schematic of the difficulty of different epilepsy etiologies that might arise in DRE patients. Since there is no biomarker for the EZ and it is never observed directly, the network mechanisms that cause seizures are complex. Case clinical complexity ordered by increasing localization difficulty: lesional (1), focal temporal (2), focal extratemporal (3), and multi-focal (4) that are present in the dataset. These four categories simplify the possible epilepsy presentations, but provide a broad categorization of simple to complex cases observed in the clinic. **(b)** Schematic of our experimental design. **(bottom row)** Shows a simplified analogous workflow that clinicians take to evaluate their confidence in a proposed *SOZ* localization resulting in a surgery. During invasive monitoring, clinicians identify the *SOZ* from iEEG patterns (e.g. spiking/rhythmic activity). When possible, subsequent surgical resection or laser ablation, generally including the *SOZ* along with a variable extent of additional tissue, is performed. Post-operatively, patients are followed for 12+ months and categorized as either success, or failure as defined in Methods - Dataset collection, resulting in an Engel or ILAE score. **(top row)** We evaluate various representations of iEEG in the form of spatiotemporal heatmaps, creating a partitioned summary of the clinically annotated *SOZ* around seizure onset, feed them into a Random Forest classifier and compute a probability of success (i.e. a confidence score) in the clinically hypothesized *SOZ*. The probability was then compared with the actual outcome of patients. These predictions can then be stratified based on clinical covariates, such as the actual surgical outcome. For a feature to be an accurate representation of the underlying epileptic phenomena, the following assumptions are made. As a result of seizure freedom, assume that the clinically hypothesized *SOZ* was sufficient, and the probability of success has a high value. In contrast, if seizures continue, then the *SOZ* was not sufficient and the probability should have a low value.

We analyze every patient’s iEEG using fragility and 20 other baseline features, resulting in spatiotemporal heatmaps for every feature. The baseline features include spectral power in various frequency bands (e.g. delta band 1-4 Hz) and specific graph measures of bivariate correlation measures (e.g. eigenvector centrality and degree of correlation and coherence matrices), which have been previously reported in the literature to correlate to the *SOZ* [11, 12, 13, 10].

### 2.1 Description of Neural Fragility

Neural fragility is a paradigm shift in the EEG analytics space. It is a concept based on the conjecture that focal seizures arise from a few fragile nodes, i.e., the SOZ, which renders the cortical epileptic network on the brink of instability. When one observes iEEG data during interictal, or preictal periods, activity recorded from each channel *appears* to hover around a baseline value (Figure 1 left). If the network is “balanced”, then it will respond transiently to an impulse, but always returns to a baseline value (e.g. Figure 2ab). In contrast, when one observes iEEG data during a seizure event, activity (i) grows in amplitude, (ii) oscillates, and (iii) spreads in the brain when the network is perturbed (Figure 1 right). This is a consequence of an “unbalanced” network that does not return to a baseline value (e.g. Figure 2c). From a dynamical systems perspective, the iEEG network has switched from a stable (non-seizure) to an unstable (seizure) network.

**Figure 2:**
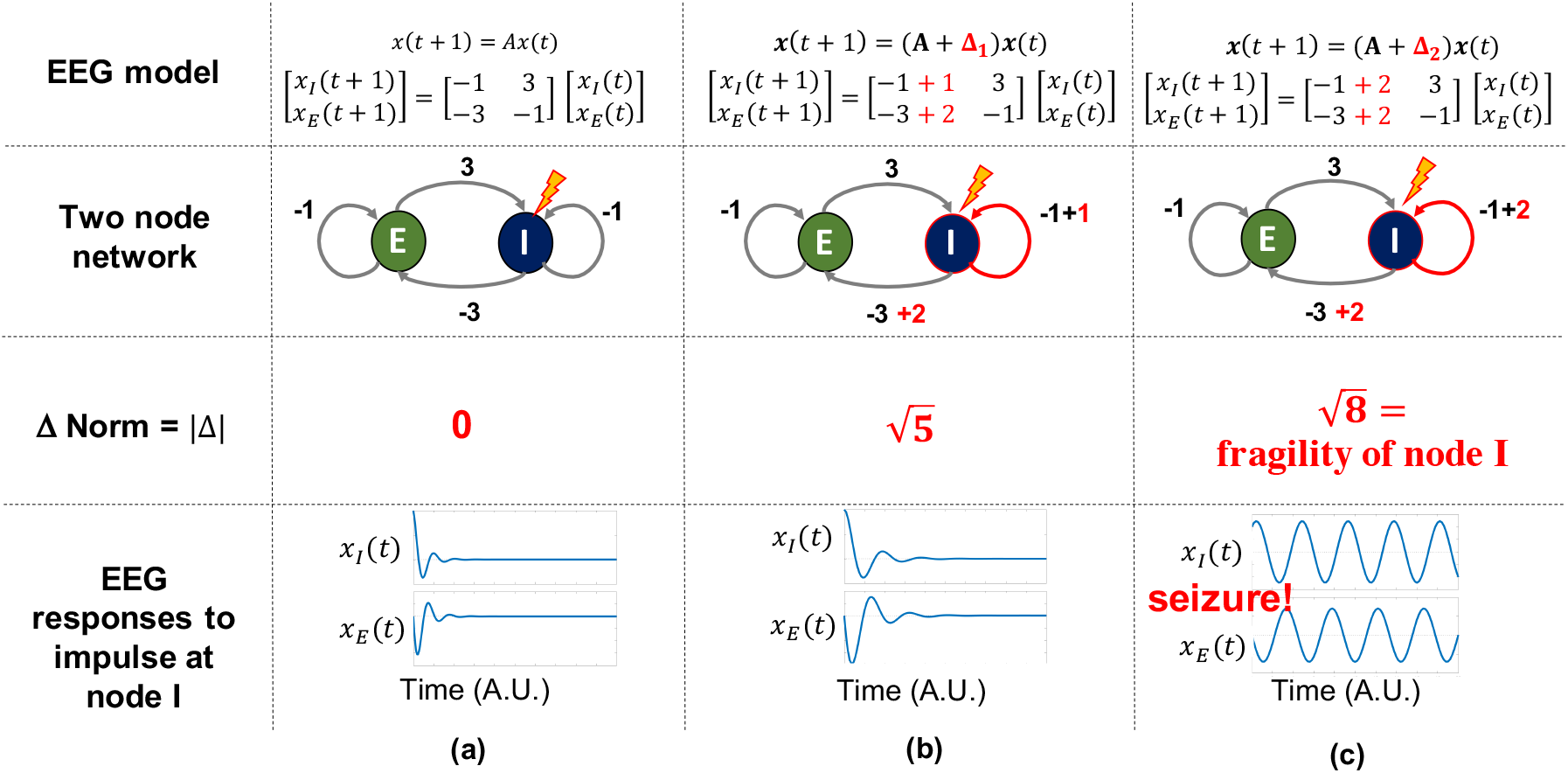
Neural fragility in a 2-node network. To build quantitative intuition on what neural fragility means in the context of a dynamical iEEG system, we construct a 2-node EEG network example with an excitatory (*E*) and inhibitory (*I*) population of neurons. For a qualitative description, see Figure 1. *x*_*I*_ (*t*) and *x*_*E*_(*t*) are the EEG activity of the *I* and E neuronal population respectively. ‘*A*’ is a linear network model quantifying how each population affects the rest over time. Δ (i.e. the fragility), is the amount of change added to a node’s connections. The fragility of the node is quantified as the minimal amount of change necessary to cause seizure-like phenomena. **(a)** shows a stable network without a perturbation added, such that the network responses due to an impulse at *I* result in a transient that reverts to baseline. **(b)** shows a perturbation added, but the network is still stable with a slightly larger transient when an impulse is applied to node *I*. Then **(c)** shows enough of a perturbation is added, such that the network becomes unstable; an impulse applied at node *I* results in oscillatory activity that does not quickly return to baseline. The magnitude of the Δ added in **(c)** is the fragility of node *I* (i.e. 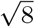).

Biologically, imbalance due to perturbations between excitatory and inhibitory connections of a neural network can occur through any number of mechanisms, such as elevated glutamate [22, 23], genetic disorder impacting synaptic inhibition [24], decreased GABA [25], inclusion of axo-axonic gap junctions [26], loss of inhibitory chandelier cells [27], or axonal sprouting from layer V excitatory pyramidal cells [28]. This imbalance within a neural network may lead to functional instability, where impulse perturbations in certain nodes lead to recurring seizures. While iEEG cannot distinguish between excitatory and inhibitory neuronal populations, the concept of imbalance causing the network to be on the brink of instability can be modeled by neural fragility at the iEEG network level.

To demonstrate how fragility is computed from a dynamical model, we consider a 2-node network as shown in Figure 2. In panel a), a stable network is shown where excitation and inhibition are balanced. The network model is provided in the top row and takes a linear form of **x**(*t* + 1) = **Ax**(*t*), where *t* is a time index (typically one millisecond). When the inhibitory node is stimulated by an impulse, both nodes transiently respond and the EEG returns to baseline (bottom row in Figure 2). In panel b), the inhibitory node’s connections are slightly perturbed in a direction that makes the inhibitory node *less* inhibitory (see red changes to its connectivity to the excitatory node). Now, when the inhibitory node is stimulated by an impulse, the responses from each node have a larger transient response but still return to baseline. Finally, in panel c), the inhibitory node’s connections are further perturbed in a direction that makes the inhibitory node *less* inhibitory. Now, when the inhibitory node is stimulated by an impulse, the responses oscillate demonstrating that the network has gone unstable. The fragility of the inhibitory node is thus quantified as 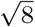 which is the norm of the perturbation vector applied to the first column in the network model. Our conjecture is that small changes in connection strengths at *SOZ* nodes cause an imbalance in connectivity between nodes in the network, resulting in susceptible seizures.

To test our conjecture, we estimate a linear time-varying model with a sliding window of *A* matrices, that characterize a linear dynamical system: *x*(*t* + 1) = *Ax*(*t*) [17, 16]. This is a generative model representing the linear dynamics between iEEG channels within a small window of time. Each A matrix is *estimated* from data via a least-squares method, and we have shown that this linear approximation is a valid model of the data [17]. From this model, we compute the minimum norm perturbation matrix, 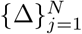, over all N channels [16] (see fig. **Computational experiment setup for all candidate** *SOZ* **features and statistical analysis** - **(a)** Any candidate feature that can produce a spatiotemporal heatmap was computed from EEG data and then partitioned by the clinically annotated *SOZ* set and the complement, *SOZ*^*C*^ (i.e. non-*SOZ* electrodes) to compute a confidence statistic measuring the feature’s belief of the clinician’s hypothesis. Here *F*_*SOZ*_ and 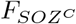 were the feature values within their respective sets. *f*_*θ*_ is the function depending on the Random Forest model parameters, *θ* that maps the statistics of the *F*_*SOZ*_ and 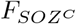 to a confidence statistic. An ideal feature would have high and low confidence for success and failed outcomes respectively. Each point on the final CS distribution comparisons represent one patient. **(b)** A more detailed schematic of how our proposed fragility and baseline features were computed from EEG data for a single snapshot of EEG data. See fragility methods section for description of x, *A* and Δ.). The 2-2-norm of each matrix represent the neural fragility for that channel. Computing the fragility now across sequentially estimated *A* matrices results in a spatiotemporal neural fragility heatmap. For a full description of neural fragility, see Methods - section Neural fragility of iEEG network.

### 2.2 Fragility highlights the clinical *SOZ* in successful patients

To qualitatively assess the usefulness of fragility in localizing electrodes of interest, we first look at specific examples of patients with varying outcomes and epilepsy etiologies (see Figure 3). We analyzed with fragility and demonstrate how it may provide additional information for SOZ localization. In Figure 4, we show an example of three different patients with differing surgical treatments, outcomes, Engel class and clinical complexity along with their fragility heatmaps and corresponding raw iEEG data (for full clinical definitions; see Supplementary Excel table).

**Figure 4:**
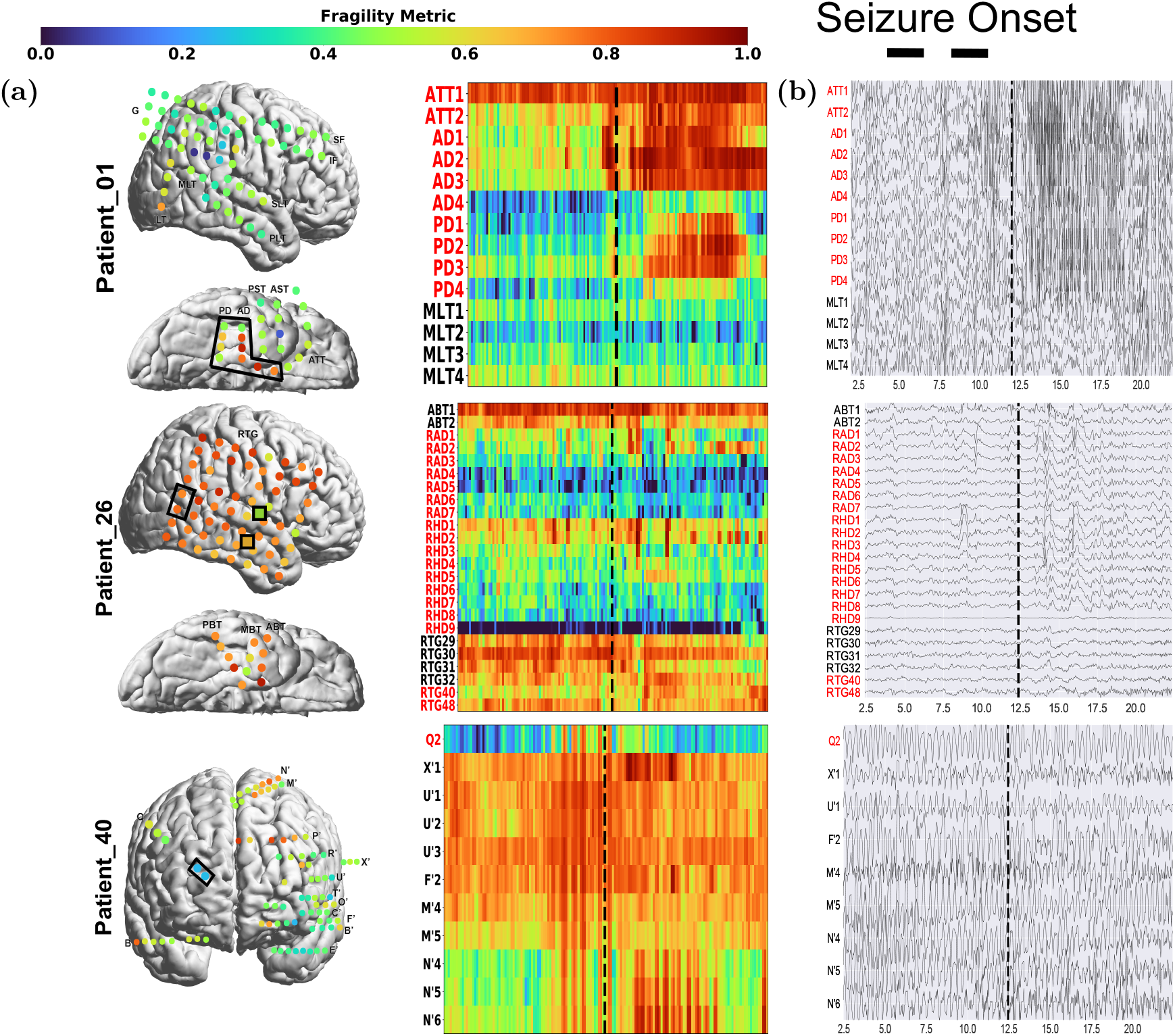
Fragility heatmaps, and corresponding raw EEG traces of successful and failed outcome patients. **(a)** From top to bottom, Patient_1 (success, NIH treated, CC1, Engel score 1), Patient_26 (failure, JHH treated, CC3, Engel score 4), and Patient_40 (failure, CClinic treated, CC4, Engel score 3) are shown respectively. The color scale represents the amplitude of the normalized fragility metric, with closer to 1 denoting fragile regions and closer to 0 denoting relatively stable regions. **(Left)** Overlaid average neural fragility value of each electrode in the window of analysis we used. Black dark squares represent a depth electrode that is not shown easily on the brain. Black lines outline where the clinicians labeled *SOZ*. Note in Patient_26, RAD and RHD electrodes are denoted by the squares with the color showing the average over the entire electrode. **(Right)** Clinically annotated heatmaps of the implanted ECoG/SEEG electrodes with red y-axis denoting SOZ contacts. The red contacts are also part of the surgical resection in these patients. Data is shown in the turbo colormap. Best seen if viewed in color. **(b)** Corresponding raw EEG data for each patient with electrodes on y-axis and time on x-axis with the dashed white-lines denoting seizure onset. Each shows 10 seconds before seizure onset marked by epileptologists, and 10 seconds after. EEG was set at a monopolar reference with line noise filtered out. Not all electrodes are visualized in the brain plot because channels that were deemed noisy, or in white matter were not included in analysis (for more information, see section Methods). In addition, only a select set of channels are chosen for the heatmap and time-series for sake of visualization on a page and to demonstrate select channels that demonstrated different fragility values. Each EEG snapshot is shown at a fixed scale for that specific snapshot that was best for visualization, ranging from 200 uV to 2000 uV.

In Figure 4a, the red electrode labels on the y-axis correspond to the clinical *SOZ* electrodes; the red electrodes are typically a subset of the resected region. It shows the period 10 seconds before and after electrographic seizure onset (black dashed line). In Patient_1, the fragility heatmap agreed with clinical visual EEG analysis, identifying the *SOZ*, which was included in the surgery and led to a seizure free patient. The clinical *SOZ* shows high degree of fragility, even before seizure onset, which is not visible in the raw EEG. The heatmap for Patient_1 also captures the propagation of seizure activity (Supplementary Figure S5). This patient had a successful surgery, and so we can assume the resected tissue likely contained the *SOZ* and EPZ regions; it is likely the clinicians correctly localized the EZ. When viewing the raw EEG data in Figure 4b (top), Patient_1 has iEEG signatures that are readily visible around seizure EEG onset (Figure 4b). We see high-frequency and synchronized spiking activity at onset that occurs in electrodes that clinicians annotated as *SOZ*, which correspond to the most fragile electrodes at onset. In addition, the fragility heatmap captures the onset in the **ATT** and **AD** electrodes and early spread of the seizure into the **PD** electrodes. Moreover, **ATT1 (anterior temporal lobe area)** shows high fragility during the entire period before seizure onset (Figure 4). This area was not identified with scalp EEG, or non-invasive neuroimaging.

### 2.3 Fragility of regions outside the *SOZ* in failed patients

In Figure 4, Patient_26 and Patient_40 both show distinct regions with high fragility that were *not* in the clinically annotated *SOZ* (or the resected region), and both had recurrent seizures after surgery. Specifically in Patient_26, the **ABT (anterior basal temporal lobe), PBT (posterior basal temporal lobe)** and **RTG29-39 (mesial temporal lobe)** electrodes were highly fragile, but not annotated as *SOZ*. Patient_26 had a resection performed in the right anterior temporal lobe region. Clinicians identified the **RAD, RHD** and **RTG40/48** electrodes as the *SOZ*. Patient_40 had laser ablation performed on the electrode region associated with **Q2**, which was not fragile. From seizure onset, many electrodes exhibit the EEG signatures that are clinically relevant, such as spiking and fast-wave activity [29]. In Patient_40, the **X’ (posterior-cingulate), U’ (posterior-insula)** and **N’/M’/F’ (superior frontal gyrus)** were all fragile compared to the **Q2** electrode, which recorded from a lesion in the right periventricular nodule. In the raw EEG data, one can see synchronized spikes and spike-waves in these electrodes, but the patient had seizures continue after resection. In the corresponding fragility heatmap, the **ABT** and the **RTG29-32** electrodes are highly fragile compared to the clinically annotated *SOZ* region. In the raw EEG shown in Figure 4b, it is not apparent that these electrodes would be part of the *SOZ*. Visual analysis of the EEG was insufficient for Patient_26 and Patient_40, which ultimately led to insufficient localizations and then to failed surgical outcomes. Based on the fragility heatmaps, the fragile regions could be hypothesized to be part of the *SOZ*, and possibly candidates for resection.

### 2.4 Fragility outperforms other features in predicting outcomes

Neural fragility in the *SOZ* was significantly higher then *SOZ*^*C*^ in successful patients as compared to failed patients (Success pvalue = 3.326e-70, Fail pvalue = 0.355; Supplementary Figure S2). Patient_01 has higher neural fragility in the *SOZ* qualitatively when compared to Patient_26 and Patient_40 (Supplementary Figure S3). This effect is seen when pooling patients across all centers as well, where neural fragility is either i) higher before the seizure onset, or ii) has a marked increase starting at seizure onset (Supplementary Figure S4). To test the validity of neural fragility and the baseline features as *SOZ* markers, we investigate each feature’s ability to predict surgical outcomes of patients when stratified by the set of *SOZ* contacts and the rest which we denote as the *SOZ* complement, *SOZ*^*C*^. We train a Structured Random Forest model (RF) for each feature that takes in a partitioned spatiotemporal heatmap based on the clinically annotated *SOZ* and generates a probability of success - a confidence score in the clinical hypothesis (for full details, see Methods Section). We test each feature’s model on a held out data set by applying a threshold to the model’s output and computing a receiver operating characteristic (ROC) curve. The ROC curve plots true positive rates versus false positive rates and the area under the curve (AUC) is a measure of predictive power of the feature. The larger the AUC, the more predictive the feature is and thus the more valid as an iEEG marker of the *SOZ*. We also compute the precision (PR) or positive predictive value (PPV) for each feature, which is the proportion of predicted successful or “positive” results that are true positives (actual successful surgeries). In addition, we compute the negative predictive value (NPV) as well. The larger the PPV/NPV, the more predictive the feature and thus the more valid it is as an iEEG marker of the *SOZ*.

Since each patient’s implantation has a varying number of electrodes, we summarize each feature’s distribution of the *SOZ* and *SOZ*^*C*^ electrodes into quantile statistics over time (for details, see Methods - Experimental design). As an input to the RF models, we compute the quantile statistics over time of the *SOZ* and *SOZ*^*C*^ set for each patient and each feature allowing each patient’s heatmaps to have the same dimensions, which are input into a machine learning RF model. RF models are attractive because they are non-parametric, are interpretable (they are a set of decision trees performing a consensus procedure), and are able to handle higher dimensional data better compared to other models, such as Logistic Regression [30]. As an output of the trained RF model, each heatmap gets mapped to a probability that the outcome will be a success. An RF model is tuned for each feature through 10-fold cross-validation (CV; see Methods-Experimental design for full details), resulting in a uniform benchmark against neural fragility on the same set of patients. The “high-gamma” frequency band feature encompasses what some would consider HFOs (i.e. 90-300 Hz).

In terms of AUC (measures discrimination), neural fragility performs the best (Figure 5a). Compared to the top 3 leading baseline features, neural fragility outperforms the beta band (15-30 Hz) power by more than 13%. The AUC of fragility is the highest with a value of 0.88 +/−0.064, compared to the next best representation, the beta frequency band, with a value of 0.82 +/−0.040. From the ROC (and PR) curves, we observe that the fragility consistently has a higher sensitivity for the same false positive rate compared to the 20 other feature representations (Supplemental Figure S6). In terms of effect size, neural fragility improves over the beta frequency band with a large effect size of 0.97 Cohen’s D with 9 out of 10 folds improving the AUC (Paired Wilcoxon Rank-sum PValue = 0.027 (see Supplemental Figure S6). As a result of the 10-fold CV, we can compare the AUC on the same set of patients in a paired effect size and statistical test improving AUC on 9 out of the 10 CV folds.

**Figure 5:**
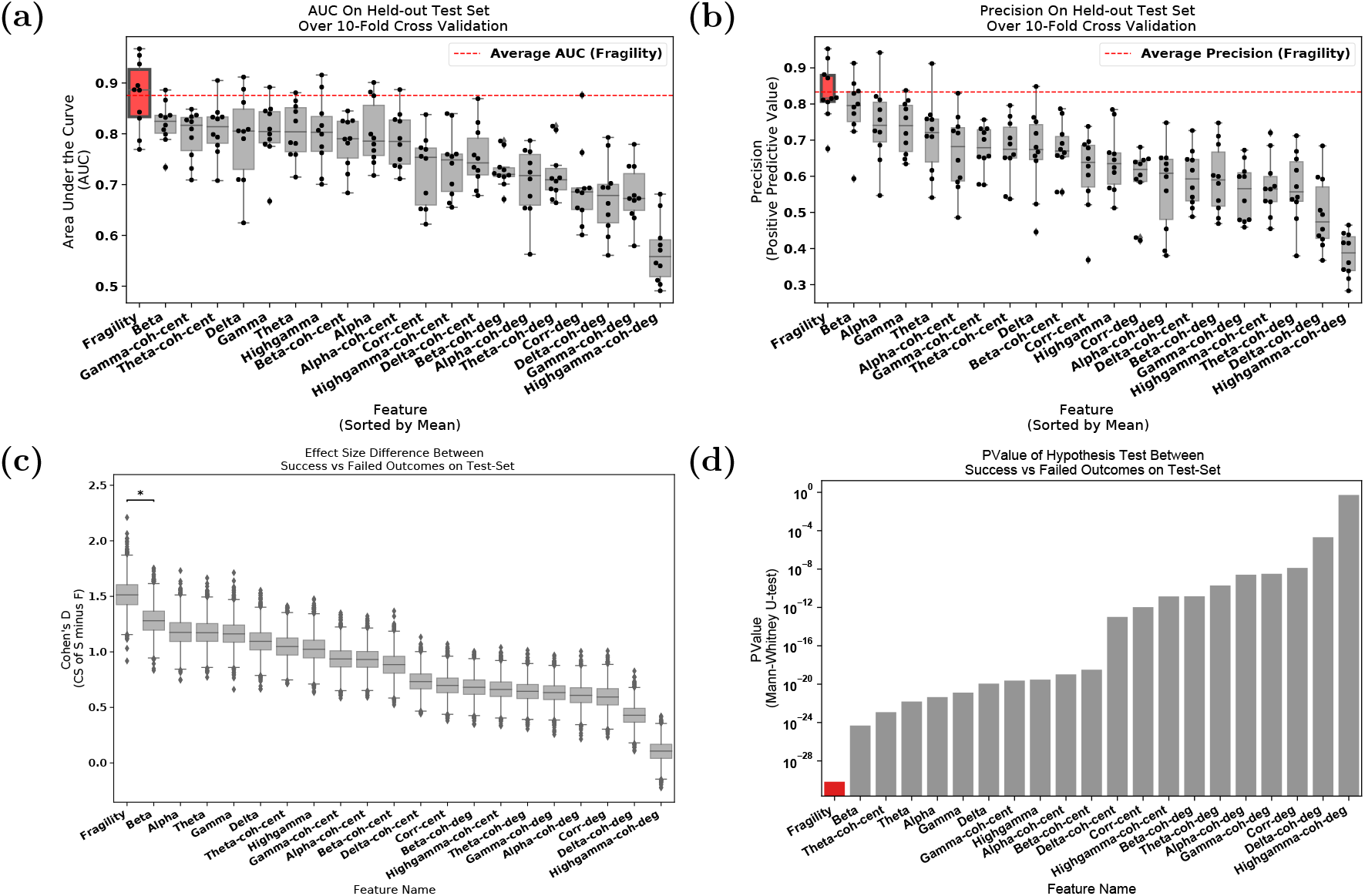
Area under the curve and average precision performance. Specific results for neural fragility are marked in red for each of the panels (a-d). **(a)** Discrimination plot (measured with AUC) shows the relative performance of benchmark feature representations compared to that achieved with neural fragility. Neural fragility has a median AUC of 0.89 (boxplot summary = 0.77, 0.97, 0.83, 0.93; min, max, first quartile, third quartile). **(b)** A similar average-PR curve shows the relative positive predictive value of all features compared with fragility. Average precision is the analagous area under the curve for the PR curve. Neural fragility has a median PR of 0.82 (boxplot summary = 0.68, 0.95, 0.81, 0.88; min, max, first quartile, third quartile). **(c)** A summary of the Cohen’s D effect size measurements between the success and failed outcome distributions across all features. The effect size of neural fragility is significantly greater then that of the beta band (alpha = 0.05). Neural fragility has a median effect size of 1.51 (boxplot summary = 0.92, 2.21, 1.43, 1.60; min, max, first quartile, third quartile). **(d)** The corresponding PValues of the effect size differences between success and failed outcomes, computed via the one-sided Mann-Whitney U-test. Note that the samples in (c) are not shown for visual sake; data was approximately bell-curve shaped and a box plot adequately summarizes the main descriptive statistics of the distribution. For box plot summary statistics (min, max, median, first quartile and third quartile) of other features, see Supplemental Data Table for this figure.

In terms of PR (measures PPV), neural fragility also performs the best (Figure 5b). IN terms of average precision, which weighs the predictive power in terms of the total number of patients, neural fragility also obtains an average precision of 0.83 +/−0.076, which is >5% better than the next best feature. In addition, compared to the clinical surgical success rate of 47%, fragility had a 76% +/−6% accuracy in predicting surgical outcome, a PPV of 0.903 +/−0.103 and a NPV of 0.872 +/−0.136. In terms of PR, neural fragility improves over the beta band power with a medium effect size of 0.57 Cohen’s D with 8 out of 10 folds improving the PR (PValue = 0.08).

When we compare the differences in the success probabilities predicted by the models stratified by surgical outcomes, we observe that neural fragility has the highest effect size of 1.507 (1.233-1.763 - 95% confidence interval) (Figure 5c) and lowest pvalue of 6.748e-31 (Figure 5d). In addition to having good discrimination, we compute how well-calibrated the model is - that is having the success probabilities values reflect true risk strata in the patient population. For example, a perfectly calibrated model would have a 20% confidence in a set of patients with exactly 20% success outcomes. We quantify how well calibrated the success probability distributions are over the held-out test set. In Supplementary Figure S6, we show that the RF model trained on neural fragility produces well-calibrated success probability values. The Brier-loss (a measure of calibration; 0 being perfect) was 0.162 +/−0.036, which was a 6.7% improvement compared to the next best feature.

### 2.5 Fragility correlates with important clinical covariates

Neural fragility correlates with treatment outcomes and clinical complexity. Successful outcomes at 12+ months post-op are defined as seizure free (Engel class I and ILAE scores of 1) and failure outcomes are defined as seizure recurrence (Engel classes 2-4). In addition, we can categorize patients by their clinical complexity (CC) as follows: (1) lesional, (2) focal temporal, (3) focal extratemporal, and (4) multi-focal (Figure 3) [21]. We stratify the distribution of the success probabilities based on Engel score and show a decreasing trend in success probability as Engel score increases (Figure 6b). The effect sizes when comparing against Engel class I were 1.067 for Engel class II (P = 4.438e-50), 1.387 for Engel class III (P = 2.148e-69), and 1.800 for Engel class IV (P = 4.476e-74). Although the AUC indirectly implies such a difference would exist between Engel score I (success) and Engel score II-IV (failures), we also observe that confidence decreases as the severity of the failure increases; Engel score IV patients experience no changes in their seizures, suggesting the true epileptic tissue was not localized and resected, while Engel score II patients experience some changes, suggesting there was portions of the true epileptic tissue that may have been resected. We also compare the success probability distributions with respect to the ILAE score, which is another stratification of the surgical outcomes (Figure 6c). There is a similar trend as seen in the Engel score - as ILAE score increases, the success probability decreases.

**Figure 6:**
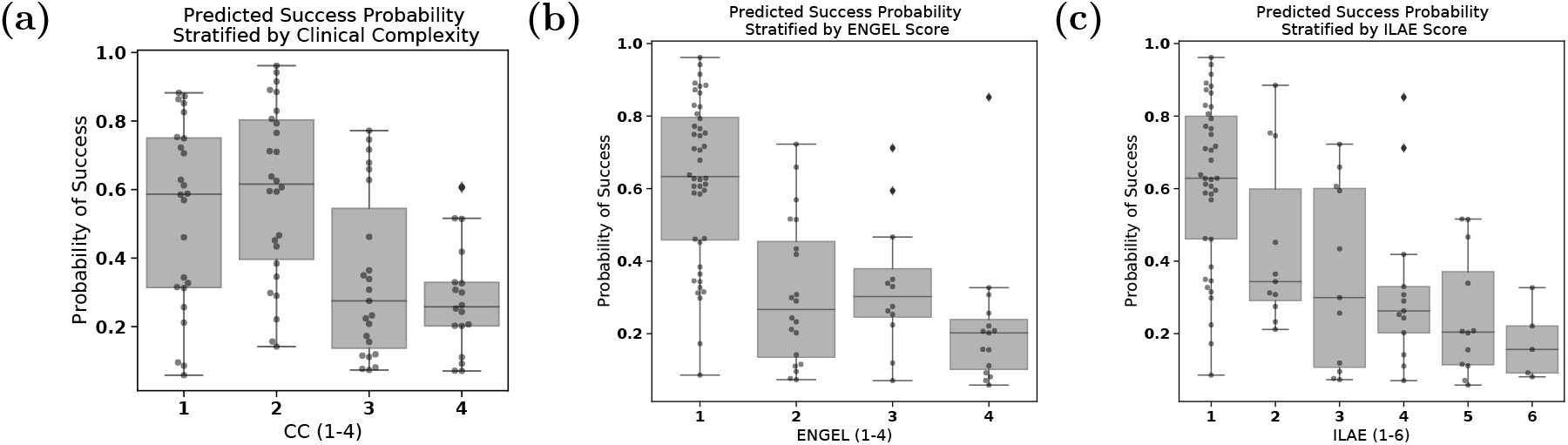
Neural fragility of patients stratified by clinical covariates. **(a)** Distribution of success probability values per patient stratified by clinical complexity (CC; see Methods-Dataset collection), where lesional (1) and temporal lobe epilepsy (2) patients have similar distributions because they are generally the “easier” patients to treat, whereas extratemporal (3) and multi-focal (4) have lower general probabilities because they are “harder” patients to treat. It is important to note that the classification experiment posed did not explicitly optimize this separation between clinical case complexities. There is a median predicted probability of success of 0.59 (boxplot summary = 0.06, 0.88, 0.31, 0.75; min, max, first quartile, third quartile) for CC 1. For CC2, CC3, and CC4, there is a median probability of success of 0.62 (boxplot summary = 0.14, 0.96, 0.40, 0.80), 0.28 (boxplot summary = 0.07, 0.77, 0.14, 0.55), and 0.26 (boxplot summary = 0.07, 0.61, 0.20, 0.33) respectively. **(b)** The distribution of the probability values per patient stratified by Engel score. Due to the AUC being high for fragility, it is expected that Engel I has high predicted probability of success, while Engel II-IV have lower success probability. However, the relative downward trend in the success probabilities from Engel II-IV indicated that neural fragility is present in the clinical *SOZ* in varying degrees from Engel II-IV, suggesting that it correlates with the underlying severity of failed outcomes. Engel IV has the lowest average predicted probability of success as expected. Engel I, II, III, and IV subjects had a median predicted probability of success of 0.63 (boxplot summary = 0.09, 0.96, 0.46, 0.80), 0.27 (boxplot summary = 0.07, 0.72, 0.14, 0.45), 0.30 (boxplot summary = 0.07, 0.71, 0.25, 0.38) and 0.20 (boxplot summary = 0.06, 0.85, 0.10, 0.24) respectively. **(c)** A similar distribution for another measure of surgical outcome, the ILAE score, where 1 are considered success and 2-6 are considered failure. Here, ILAE 2-5 follow a decreasing trend with ILAE-6 having the lowest average predicted probability of success. ILAE 1-6 has a median predicted probability of success of 0.63 (boxplot summary = 0.09, 096, 0.46, 0.80), 0.34 (boxplot summary = 0.21, 0.88, 0.29, 0.60), 0.30 (boxplot summary = 0.07, 0.72, 0.11, 0.60), 0.26 (boxplot summary = 0.07, 0.85, 0.20, 0.33), 0.20 (boxplot summary = 0.06, 0.52, 0.11, 0.37), and 0.16 (boxplot summary = 0.08, 0.33, 0.09, 0.22) respectively.

We also analyze the success probability with respect to the epilepsy severity measured by the clinical complexity (CC) of the patient, which is a categorization of the etiology of the disease we used. CC is determined by what type of seizures the patient exhibits rather than the severity of the seizures after-surgery (for more details see Methods - Dataset collection). CC is a factor that is determined before surgery and one we expect would correlate with failure rate. The higher the CC, the more difficult localization of the *SOZ* is and hence possibly the more seizure recurrences after surgery. In Figure 6a, we observed that CC1 and CC2 (i.e. lesional and temporal lobe) patients had a similar distribution of success probabilities (Cohen’s D = 0.020; MW U-test PValue = 0.799; LQRT PValue = 0.76), while CC3 and CC4 had significantly lower distributions (CC3: [Cohen’s D = 0.779; MW U-test PValue = 2.808e-19; LQRT PValue = 0.00]; CC4: [Cohen’s D = 1.138; MW U-test PValue = 7.388e-26; LQRT PValue = 0.00]). This trend is not optimized for directly in our discrimination task, but it aligns with clinical outcomes. CC1 and CC2 are comparable, which agrees with current data suggesting that lesional and temporal lobe epilepsy have the highest rates of surgical success [31, 32]. Extratemporal (CC3) and multi-focal (CC4) patients tend to have lower success rates due to insufficient localizations and thus the neural fragility confidence in those *SOZ* localizations should be low [31].

The highest values of neural fragility (i.e. red zones) that differentiates the *SOZ* and *SOZ*^*C*^ are what contribute most to making a correct classification (Supplemental Figure S7). However, in general neural fragility is not correlated with any single frequency band (Supplemental Figure S8). We also show how fragility heatmaps are the most interpretable over all baseline features. When comparing the contrast between *SOZ* and *SOZ*^*C*^ of success and failed patients, neural fragility has the largest difference of (Supplemental Figure S9). Finally, we examine the success probability differences based on gender, handedness, onset age and surgery age to show that there are no relevant differences (Supplemental Figure S10).

## 3 Discussion

We demonstrate that neural fragility, a networked-dynamical systems based representation of iEEG is a strong candidate for a highly interpretable iEEG biomarker of the *SOZ*. We compared neural fragility to 20 other popular features using data from n=91 patients treated across five centers with varying clinical complexity. Neural fragility performed the best in terms of AUC, PR and interpretability, bench-marking proposed iEEG features in a uniform fashion over the same sets of patients using the best possible parameters of a RF model to compute a confidence measure (i.e. probability of success) of the clinically annotated *SOZ*. We spent over four years to successfully collect and annotate this heterogeneous dataset. To facilitate reproducibility and further investigation of iEEG features, we made this dataset BIDS-compliant and publicly available through the OpenNeuro website (for details see Methods - Code availability).

### Challenges in validating iEEG features as SOZ markers

Many features have been proposed as potential biomarkers for the SOZ, but none have successfully translated into the clinical workflow [11, 12, 33, 31]. Current limitations for evaluating computational approaches to localization are largely attributed to i) insufficient benchmarking to other iEEG features and ii) a lack in sufficient sampling across epilepsy etiologies.

Even with a seemingly successful feature derived from data, it is important to benchmark against existing approaches to provide a holistic view of the value of the said feature. Without benchmarking, it is easy to become overly optimistic in terms of the performance of a feature, whereas it may be that other iEEG features perform just as well due to the low clinical complexity presentation of the patient. While other traditional features such as the power in the beta band seem to be informative in *SOZ* localization, neural fragility outperforms in effect size, p-value and interpretability.

Although predicting surgical outcomes in our experimental setup is promising, it will be important to understand why certain localizations are successful and why certain are insufficient. For example, in patients with lesions on MRI scans that correlate with the patient’s EEG and seizure semiology, surgical resection can lead to seizure freedom in approximately 70% of patients [21]. Even in these relatively straightforward cases, localization is not perfect, possibly due to chronic effects of epilepsy such as kindling, which can cause neighboring tissue to become abnormal and epileptogenic [34]. There is a need to sample a heterogeneous and large patient population and derive a feature is invariant to epilepsy type and clinical covariates.

### Clinical case complexity and surgical outcomes

In our dataset, we saw varying outcomes across the five clinical centers. Difference in seizure outcomes across the clinical centers can be explained by several contributory factors affecting the complexity of the epileptic syndromes. The multi-factorial contributory factors can be related to i) the percentage of non-lesional versus lesional cases, ii) multiple surgical interventions in the past, iii) patient selection and iv) group experience. For example, JHH is a tertiary referral center with a great deal of experience, but also high complexity cases (non-lesional MRIs, multiples surgical interventions in the past, complex semiology and EEG interpretation). In addition, data collection is limited depending on clinician resources and retrospective data availability. The relative center-by-center outcomes are not reflective of the actual center’s surgical outcome rate, but rather samples from the clinical cases that those clinicians analyzed and data were readily available for sharing. The pooled 91 patients though do reflect the approximately 50% surgical success rates seen in DRE patients [20, 35, 36]. By including a multitude of centers, we sought to build a diversified sample of varying clinical complexities and practices, thus lending the evaluation of neural fragility more confidence. An important next step would be the prospective evaluation of neural fragility in the context of different clinical case complexities to determine if surgical treatment can be improved with *SOZ* localization assistance. In addition, as much of the dataset should be made BIDS-compliant and available to the public as possible to further developments of localization algorithms.

### Limitations of the most popular iEEG features

A google scholar search using the keywords: “localization of seizure onset zone epilepsy intracranial EEG” produces over 24,000 results. This is striking considering no computational tools to assist in SOZ localization are in the clinical workflow today. The majority of proposed features lack consistency in finding objective iEEG quantities that correlate to clinically annotated SOZ because they fail to capture internal properties of the iEEG network which are critical to understand when localizing the SOZ. Proposed algorithms either (i) compute EEG features from individual channels (e.g. spectral power in a given frequency band), thus ignoring dependencies between channels [9, 37] to name a few, or they (ii) apply network-based measures to capture pairwise dependencies in the EEG window of interest [11, 12]. Specifically, correlation or coherence between each pair of EEG channels is computed and organized into an adjacency matrix, on which summary statistics are derived including degree distribution and variants of centrality [11, 12, 13, 10, 15]. Such network-based measures are not based on well formulated hypotheses of the role of the epileptic tissue in the iEEG network, and many different networks (adjacency matrices) can have identical summary statistics resulting in ambiguous interpretations of such measures [38].

A popular EEG feature that has been proposed as an iEEG marker of the SOZ and reported in over 1000 published studies is High-frequency oscillations (HFOs) ([9, 39, 33] to name a few). HFOs are spontaneous events occurring on individual EEG channels that distinctively stand out from the background signal and are divided into three categories: ripples (80–250 Hz), fast ripples (250–500 Hz), and very-fast ripples (>500 Hz) [40]. Retrospective studies suggested that resected brain regions that generate high rates of HFOs may lead to good post-surgical outcome (e.g., [41, 42]). Although they found significant effects for resected areas that either presented a high number of ripples or fast ripples, effect sizes were small and only a few studies fulfilled their selection criteria [42]. Furthermore, several studies have also questioned the reproducibility and reliability of HFOs as a marker [43, 41, 39] and there are also physiologic, non-epileptic HFOs. Their existence poses a challenge, as disentangling them from clinically relevant pathological HFOs still is an unsolved issue [44].

Similar inconclusive results hold in completed prospective studies of HFOs. In 2017, an updated Cochrane review [45] investigated the clinical value of HFOs regarding decision making in epilepsy surgery. They identified only two prospective studies at the time and concluded that there is not enough evidence so far to allow for any reliable conclusions regarding the clinical value of HFOs as a marker for the SOZ. Today, five clinical trials are listed as using HFOs for surgical planning on *clinicaltrials*.*gov* as either recruiting, enrolling by invitation, or active and not enrolling and none have reported results. The fundamental limitation of the aforementioned studies lies in the fact that they approach the SOZ EEG marker discovery process as a signal processing and pattern recognition problem, concentrating on processing EEG observations to find events of interest (e.g. HFOs) as opposed to understanding how the observations were generated in the first place and how specific internal network properties can trigger seizures.

### Why Neural Fragility Performs Well

Rather than analyzing iEEG data at the channel and signal processing level, we seek to model the underlying network dynamics that might give rise to seizures in the form of neural fragility in a dynamical network model. A notion of fragility in networks is commonly seen in analysis of structural [46] network. Although we are not directly analyzing the structural nature of neuronal network, there are studies that have characterized epilepsy in terms of structural fragility and network organization [47, 18]. Specifically, in cellular studies [28, 27], epilepsy is caused by changes in the structural cell network, which causes loss of inhibition or excessive excitation; these biological changes cause downstream aberrant electrical firing (i.e. seizures). By quantifying the fragility of each node in an iEEG network, we determine how much of a change in that region’s dynamical connections is necessary to cause seizure-like phenomena (e.g. instability). As a result, high neural fragility coincides with a region that is sensitive to minute perturbations, causing unstable phenomena in the entire network (i.e. a seizure).

### Scientific Advances Emerging from Neural Fragility

Neural fragility has the potential to re-define how epilepsy surgery is performed, departing from the classical “localization paradigms” and “en-bloc resections” to a personalized “network-based” user-friendly visualization and surgical strategy. By developing a 3D (brain region, time, fragility) network-based method for anatomical representation of the epileptiform activity, including the seizure onset areas and the early propagation zone, this study will have the potential to offer a safer, more efficient, and cost-effective treatment option for a highly challenging group of patients with disabling DRE. More precise SOZ localization using neural fragility would also guide of chronic implantation of neurostimulation devices aimed to suppress seizures with bursts of current when detected [48, 49]. Neural fragility may also be relevant in detecting epileptogenic regions in the brain when applied to interictal (between seizures) iEEG recordings. Seizure iEEG data are currently the gold standard in clinical practice for localizing the *SOZ* [21]. However, having patients with electrodes implanted for long periods of time, and requiring the monitoring of multiple seizure events over many weeks carries the risk of infection, sudden death, trauma and cognitive deficits from having repeated seizures. This contributes to the large cost of epilepsy monitoring [2]. If a candidate iEEG marker could be found that is able to provide strong localizing evidence using only interictal data, then it would significantly reduce invasive monitoring time [47].

Neural fragility is an EEG marker that can also further advance our knowledge of neural mechanisms of seizure generation that will then drive more effective interventions. For example, fragility can be used to identify pathological tissue that are then removed and tested in vitro for abnormal histopathological structure [28, 27]. Knowledge of structural abnormalities may inform new targeted drug treatments. In the future, specific fragility patterns can be correlated with specific pathological substrates. Likely, the specific pathological substrates will have different therapeutic approaches. As an example, epilepsy caused by focal cortical dysplasia is treated with focal surgical resection, but post-encephalitic epilepsy may have a better therapeutic response with immunosuppressants and steroids.

Finally, neural fragility may have broader implications in understanding how underlying brain network dynamics change during intervention (e.g. drugs or electrical stimulation). Fragility analysis can be applied as a method of assessing the efficacy of specific drug trials to specific pathological groups, which include not only epilepsy but other neurological conditions such as Alzheimer’s’s disease or the spectrum of dementias. Commonly, the current optimal criteria to recognize therapeutic success in many neurological conditions is purely clinical, but clinical responses are not immediate. This delay in recognizing the appropriate drug and adequate therapeutic doses is highly detrimental. Computational methods as fragility measuring EEG before and after drug administration could provide an additional criteria for drug responses. This quantitative measurement can be immediate, guiding the treating physician to the correct treatment without delays and unnecessary drug trials. Furthermore, if neural fragility could be accurately obtained from non-invasive tests or from permanently implanted devices, the current fragility of the network could be used as a surrogate marker of patient’s current clinical state. As such, the changes in the fragility could be used as a proxy for improvement or recurrences that occur as medication doses (or other treatments, such as Keto Diet [50]) are changed over time.

## Supporting information

Patient de-identified metadata.

Supplemental box plot summary for figure 5a

Supplemental box plot summary for figure 5b

Supplemental box plot summary for figure 5c

Supplemental box plot summary for figure 6a

Supplemental box plot summary for figure 6b

Supplemental box plot summary for figure 6c

## 4 Acknowledgements

AL is supported by NIH T32 EB003383, the NSF GRFP (DGE-1746891), the Arcs Chapter Scholarship, Whitaker Fellowship and the Chateaubriand Fellowship. SVS is supported by NIH R21 NS103113, the Coulter Foundation, Maryland Innovation Initiative, US NSF Career Award 1055560 and the Burroughs Wellcome Fund CASI Award 1007274. EJ is supported by NIA K23 AG063899. IC was partially supported by NIH R25NS108937-02. KZ and SI are supported by the Intramural Research Program of the National Institute of Neurological Disorders and Stroke. SC is supported by 1UL1TR003098-01 and the UMB Institute for Clinical and Translational Research. JHopp is supported by NINDS 1 UO1 NS08803401 - Established Status Epilepticus Treatment Trial (ESETT), MII TA Grant 20180828 TEDCO - EpiWatch: A Mobile Health Application for Epilepsy Management and U10 NS058932-01 NIH NINDS - “Neurological Emergencies Treatment Trials (NETT) Network Clinical/Site Hub”. Computational resources were also provided by the Maryland Advanced Research Computing Center (MARCC). The authors would like to thank Carey Priebe for useful discussions on statistical analyses, Joshua Vogelstein for introducing AL to the concepts of Structured Random Forests, Sarah Kim and Rachel June Smith for very helpful reviews of the manuscript and Macauley Breault for help in creating brain figures. Finally, the authors would like to thank the editor and anonymous reviewers for their insightful comments and reviews.

## 5 Author Contributions Statement

AL, JGM and SVS conceived the project. AL, IC, DB, JJ, AC, AK, JHopp, SC, JHaagensen, EJ, WA, NC, ZF, JB and JM collected and supervised human epilepsy recordings and processes related to IRB at their respective centers. IC, DB, JJ, AC, AK oversaw data from University of Miami Hospital. JHopp, SC, and JHaagensen oversaw data from University of Maryland Medical Center. EJ, WA and NC oversaw data from Johns Hopkins Hospital. SI and KZ oversaw data from National Institute of Health. ZF, JB and JGM oversaw data from Cleveland Clinic. AL organized the data from all centers, converted data to BIDS, conducted the analyses and generated the figures. AL and SVS wrote the paper with input from the other authors. CH helped develop the structured random forest model and statistical analyses.

## 6 Competing Interests Statement

The authors declare the following competing interests: AL, SVS and JM have equity in a startup, Neurologic Solutions Co., related to epilepsy data analysis. WA is on the Board of Advisors of Longeviti Neuro Solutions and a Compensated Consultant at Globus Medical. JHopp reports the following disclosures: Personal compensation: Royalties from publishing – UpToDate, Inc., Honoraria from speaking engagements - J. Kiffin Penry Epilepsy Education Programs, Expert witness consultation: Gerolamo McNulty Divis & Lewbart A Professional Corporation Attorneys At Law Goodell, Devries, Leech & Dann, LLP. All other authors declare no competing interests.

**Table 1:** A table of number of patients per clinical center (NIH = National Institute of Health; JHH = Johns Hopkins Hospital; UMMC = University of Maryland Medical Center; UMH = University of Miami Jackson Memorial Hospital; CClinic = Cleveland Clinic). Includes number of success, failures, and gender per group. Note that JHH did not retain gender information for these group of patients.

| Clinical Center Patients |  |  |  |  |  |  |
| --- | --- | --- | --- | --- | --- | --- |
| Clinical Covariate | NIH | JHH | UMMC | UMH | CClinic | Total |
| Number Patients | 14 | 4 | 7 | 5 | 61 | 91 |
| Number Success | 9 | 1 | 7 | 3 | 24 | 44 |
| Number Failure | 5 | 3 | 0 | 2 | 37 | 47 |
| Number Male | 6 | n/a | 7 | 1 | 30 | 44 |
| Number Female | 8 | n/a | 0 | 4 | 31 | 43 |

## 8 Methods

All data were acquired with approval of local Institutional Review Board (IRB) at each clinical institution: UMMC by IRB of the University of Maryland School of Medicine, UMH by University of Miami Human Subject Research Office - Medical Sciences IRB, NIH by the National Institute of Health IRB, JHH by Johns Hopkins IRB and CClinic by Cleveland Clinic Institutional Review Board. Informed consent was given at each clinical center. The acquisition of data for research purposes was completed with no impact on the clinical objectives of the patient stay. Digitized data were stored in an IRB-approved database compliant with Health Insurance Portability and Accountability Act (HIPAA) regulations.

### 8.1 Data availability

We also released the raw iEEG data for patients from NIH, UMH, UMMC, and JHH in the Open-Neuro repository in the form of BIDS-iEEG (https://openneuro.org/datasets/ds003029). Due to restrictions on data sharing from CClinic, we were unable to release the raw iEEG data that we received from this site. Dataset from CClinic is available upon request from authors at the CClinic.

### 8.2 Code availability

All main figures of the manuscript can be reproduced using Gigantum at https://gigantum.com/adam2392/neural-fragility-ictal-study-figures. We include a jupyter notebook written in Python to help reproduce figures. An example of neural fragility being run on Patient_01 can be shared upon request due to licensing restrictions.

The neural fragility algorithm has been implemented in a FDA 510k approved software medical device (current 510k number: K201910). More information will be available at www.neurologicsolutions.co. Otherwise, please contact corresponding authors for more information regarding a clinical demonstration.

### 8.3 Dataset collection

iEEG data from 91 DRE patients who underwent intracranial EEG monitoring, which included either electrocorticography (ECoG), or depth electrodes with stereo-EEG (SEEG) were selected from University of Maryland Medical Center (UMMC), University of Miami Jackson Memorial Hospital (UMH), National Institute of Health (NIH), Johns Hopkins Hospital (JHH), and the Cleveland Clinic (CClinic). Patients exhibiting the following criteria were excluded: no seizures recorded, pregnant, sampling rate less than 250 Hz, previous surgeries more then 6 months before, and no surgery performed (possibly from *SOZ* localizations in eloquent areas). We define successful outcomes as seizure free (Engel class I and ILAE scores of 1 and 2) at 12+ months post-op and failure outcomes with seizure recurrence (Engel classes 2-4) [1, 2, 3]. Of these 91 patients, 44 experienced successful outcomes and 47 had failed outcomes (age at surgery = 31.52 +/−12.32 years) with a total of 462 seizures (seizure length = 97.82 +/−91.32 seconds) and 14703 total number of recording electrodes (159.82 +/−45.42 per subject). Decisions regarding the need for invasive monitoring and the placement of electrode arrays were made independently of this work and part of routine clinical care. Data collection and labeling of patients were done retrospectively and thus blind to the authors conducting data analysis (i.e. AL, CH, SVS), but not with respect to the corresponding clinicans. Data analysis were not performed blind to the conditions of the experiments. Sample sizes were determined by attempting to gather as many patients as possible from multiple clinical centers. When possible, we sought to have a balance of lesional, temporal, extra-temporal and multi-focal epilepsy patients, along with a relatively proportional number of success and failed epilepsy surgery patients (success = seizure freedom, failure = seizure recurrence). We collected 100 patients and then removed 9 due exclusion criterion. No statistical methods were used to pre-determine sample sizes but our sample sizes are larger then those reported in previous publications [4].

Our clinician team categorized patients by surgical outcome, Engel class and ILAE score. In addition, we categorized patients by their clinical complexity (CC) as follows: (1) lesional, (2) focal temporal, (3) focal extratemporal, and (4) multi-focal (Figure 3) [3, 5]. Each of these were categorized based on previous outcome studies that support this increasing level of localization difficulty. Lesional patients have success rates of 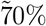, experiencing the highest rate of surgical success because the lesions identified through MRI are likely to be part of the SOZ [6]. Localization and surgical success are more challenging patients with non-lesional MRI, with average surgical success rates in temporal, extratemporal and multi-focal epilepsy of 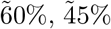 and 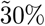, respectively [7, 8]. Patients that fit into multiple categories were placed into the more complex category. Next, clinicians identified electrodes that they hypothesized as *SOZ*. In general, this was a subset of the resected region for all patients, unless otherwise noted. The epileptologists define the clinically annotated *SOZ* as the electrodes that participated the earliest in seizures. Every patient’s *SOZ* was labeled by 1-3 trained epileptologists (depending on the center). The corresponding *SOZ* complement that we define, or *SOZ*^*C*^, are the electrodes that are not part of the *SOZ*. For more detailed information regarding each patient, see Supplemental clinical data summary Excel file and A table of number of patients per clinical center (NIH = National Institute of Health; JHH = Johns Hopkins Hospital; UMMC = University of Maryland Medical Center; UMH = University of Miami Jackson Memorial Hospital; CClinic = Cleveland Clinic). Includes number of success, failures, and gender per group. Note that JHH did not retain gender information for these group of patients..

At all centers, data were recorded using either a Nihon Kohden (Tokyo, Japan) or Natus (Pleasanton, USA) acquisition system with a typical sampling rate of 1000 Hz (for details regarding sampling rate per patient, see Supplementary file table). Signals were referenced to a common electrode placed subcutaneously on the scalp, on the mastoid process, or on the subdural grid. At all centers, as part of routine clinical care, up to three board-certified epileptologists marked the electrographic onset and the termination of each seizure by consensus. The time of seizure onset was indicated by a variety of stereotypical electrographic features which included, but were not limited to, the onset of fast rhythmic activity, an isolated spike or spike-and-wave complex followed by rhythmic activity, or an electrodecremental response. Each snapshot available for a patient was clipped at least 30 seconds before and after the ictal event. We discarded electrodes from further analysis if they were deemed excessively noisy by clinicians, recording from white matter, or were not EEG related (e.g. reference, EKG, or not attached to the brain) which resulted in 97.23 +/−34.87 (mean +/−std) electrodes used per patient in our analysis. Data was initially stored in the form of the European Data Format (EDF) files [9]. We preprocessed data into the BIDS-iEEG format and performed processing using Python3.6, Numpy, Scipy, MNE-Python and MNE-BIDS [10, 11, 12, 13, 14, 15, 16]. Figures were generated using Matplotlib and Seaborn [17, 18, 19]. Statistical analyses were performed with mlxtend, pingouin, dabest, scipy and scikit-learn [20, 21, 22, 16, 23].

### 8.4 Preprocessing of data

Every dataset was notch filtered at 60 Hz (with a cutoff window of 2 Hz) and bandpass filtered between 0.5 and the Nyquist frequency with a fourth order Butterworth filter. A common average reference was applied to remove any correlated noise. EEG sequences were broken down into sequential windows and the features were computed within each window (see Methods - Neural fragility of iEEG network, Baseline features - spectral features and Baseline features - graph analysis of networks for details). Each proposed feature representation produces a value for each electrode for each separate window, and results in a full spatiotemporal heatmap when computed over sequential windows with electrodes on the y-axis, time on the x-axis and feature value on the color axis. In total, we computed 20 different baseline feature representations from the iEEG data: 6 frequency power bands, 7 eigenvector centralities (one for each frequency band coherence connectivity matrix and one for a correlation connectivity matrix), 7 degrees (one for each frequency band coherence connectivity matrix and one for a correlation connectivity matrix). Values at each window of time were normalized across electrodes to values that could range from 0 up to at most 1, to allow for comparison of relative feature value differences across electrodes over time; the higher a normalized feature, the more we hypothesized that electrode was part of the SOZ [24].

### 8.5 Neural fragility of iEEG network

When one observes iEEG data during interictal, or preictal periods, activity recorded from each channel is noisy and hovers around a baseline value. In contrast, when one observes iEEG data during a seizure event, activity (i) grows in amplitude, (ii) oscillates, and (iii) spreads in the brain. From a dynamical systems perspective, the iEEG network has switched from a stable (non-seizure) to an unstable (seizure) network. The only difference between the iEEG networks in Figure 2 is the connection strengths representing the dynamical interactions between a few channels, i.e., the *SOZ*. Our conjecture is that small changes in connection strengths at *SOZ* nodes cause an imbalance in inhibitory and excitatory connectivity between brain regions. Either inhibition is decreased and/or excitation is increased; thus, if the *SOZ* is perturbed then over excitation can occur manifesting in a seizure.

To compute fragility heatmaps from iEEG recordings, we first constructed simple linear models as described above and in equation 1.

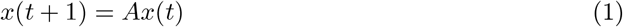

Since each observation, x ∈ ℝ^*d*^, has dimension d (number of channels), we would like to formulate a least-squares estimation procedure with *n > d* samples. We choose n to represent a 250 ms iEEG window. We then have the following representation of *X*(*t*) ∈ R^*d×n* − 1^:

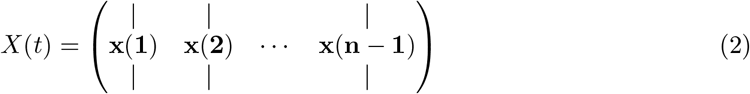

and the following representation for *X*(*t* + 1) ∈ ℝ^*d*×*n* − 1^

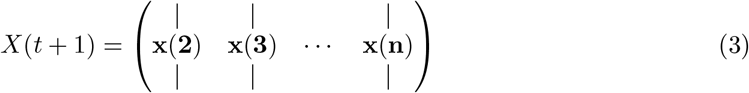

The least-squares will now seek to fit a linear operator *A* such that:

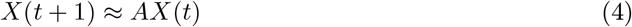

This linear operator representation of the dynamical system has connections to Koopman operator theory [25] and Dynamic Mode Decomposition in Fluid Mechanics [26]. We seek to approximate the inherently nonlinear iEEG dynamics within a small window of time using a finite-dimensional approximation of the Koopman operator using the observables (i.e. *x*(*t*)) themselves. We specifically used least-squares algorithm with a 10e-5 l2-norm regularization to ensure that the model identified was stable (with absolute value of eigenvalues ≤ 1) as in [27, 24]. Then, we slid the window 125 ms and repeated the process over the entire data sample, generating a sequence of linear network models in time as in Supplemental Figure S1b).

We systematically computed the minimum perturbation required for each electrode’s connections (Supplemental Figure S1b) to produce instability for the entire network as described in [24]. This is represented in equation 5, where the Δ_*i*_ is the desired column perturbation matrix for channel i, and the *λ* = *r* ∈ ℂ is the desired radii to perturb one single eigenvalue to.

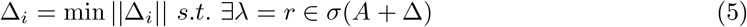

More specifically, we compute a structured perturbation matrix, such that:

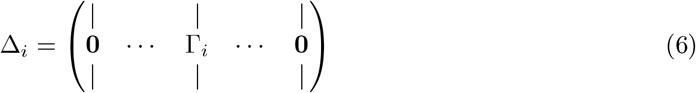

where each Γ_*i*_ ∈ ℝ^*d*^ is the actual column perturbation vector. The intuition for using this type of structured perturbation is described in the main paper in section Description of Neural Fragility. We demonstrate how to solve for this using least-squares in [24]. The electrodes that were the most fragile were hypothesized to be related to the SOZ in these epilepsy networks (seen as the dark-red color in the turbo colormap in Figure 4).

### 8.6 Baseline features - spectral features

We constructed spectral-based features from frequency bands of interest by applying a multi-taper Fourier transform over sliding windows of data with a window/step size of 2.5/0.5 seconds [28]. We required relatively longer time windows to accurately estimate some of the lower frequency bands. Each EEG time series was first transformed into a 3-dimensional array (electrodes X frequency X time), and then averaged within each frequency band to form six different spectral feature representations of the data. We break down frequency bands as follows:

1. Delta Frequency Band [0.5 - 4 Hz]
2. Theta Frequency Band [4 - 8 Hz]
3. Alpha Frequency Band [8 - 13 Hz]
4. Beta Frequency Band [13 - 30 Hz]
5. Gamma Frequency Band [30 - 90 Hz]
6. High-Gamma Frequency Band [90 - 300 Hz]

The High-Gamma frequency band includes frequencies of “ripples” - a category of HFOs.

### 8.7 Baseline features - graph analysis of networks

We computed a time domain model using Pearson correlation (equation 7) and a frequency domain model using coherence (equation 8). We computed the connectivity matrix using MNE-Python and used the default values [13]. In the equations, (*i, j*) are the electrode indices, Cov is the covariance, *σ* is the standard deviation, *f* is the frequency band, and *G* is cross-spectral density. These connectivity models attempt to capture linear correlations either in time, or in a specific frequency band, but are not dynamical system representations of the data (i.e. *x*(*t* + 1) = *Ax*(*t*)). For each network-based feature, a sliding window/step size of 2.5/0.5 seconds were used, resulting in a sequence of network matrices over time resulting in 3-dimensional arrays (electrodes X electrodes X time) [4, 28, 29].

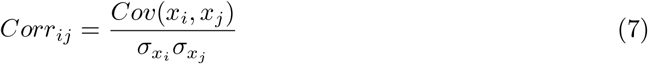

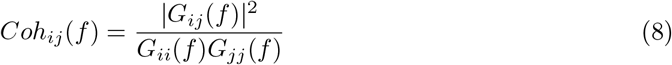

From each network matrix, we computed the eigenvector centrality [28, 4], and the degree features of the network for each electrode across time. Since coherence and pearson correlation are symmetric matrices, the in-degree is equivalent to the out-degree. Centrality describes how influential a node is within a graph network. The degree is the weighted sum of the connections that connect to a specific node. Both features are potential measures that attempt to capture the influence of a specific electrode within an iEEG network, through the lens of graph theory.

### 8.8 Experimental design

We tested if the neural fragility representation of iEEG data localized the *SOZ* better compared to other proposed features, compared to clinicians and compared to chance. Electrodes with extreme feature values deviating from the average were hypothesized as part of the *SOZ*.

To compute a probability of successful surgical outcome for each patient, we trained a non-parametric machine learning classifier, a Random Forest classifier to output a probability value that the surgery was a success. We input the distribution of feature values in the clinically *SOZ* and the *SOZ*^*C*^. We considered a set of hyperparameters that we then performed evaluation over a 10-fold nested cross-validation scheme. We then performed statistical analysis on the final classification performance to determine the most robust feature representation.

#### Pooled Patient Analysis

First, we analyzed the difference in the distributions of neural fragility between *SOZ* and the *SOZ*^*C*^. We pooled all patients together, stratified by surgical outcome and compared the neural fragility distributions using a one-sided Mann-Whitney U test (Success pvalue = 3.326e-70, Fail pvalue = 0.355; Supplementary Figure S2). This suggested that there is some effect on average where fragility is higher in the *SOZ* for successful outcomes, so we next looked at the distributions per patient’s seizure snapshot around seizure onset. In Supplementary Figure S3, success outcome patients have a higher neural fragility in the *SOZ*. This effect is seen when pooling patients across all centers as well, where neural fragility is either i) higher before the seizure onset, or ii) has a marked difference starting at seizure onset (Supplementary Figure S4). Next, we performed a classification experiment (Figure 3) that would determine the robustness of the neural fragility representation at the patient level benchmarked against 20 other features.

#### Non-parametric Decision-Tree Classifier

To determine the value of a feature representation of the iEEG data, we posed a binary classification problem where the goal would be to determine the surgical outcome (success or failure) for a particular patient’s spatiotemporal heatmap. Each spatiotemporal heatmap was split into its *SOZ* and *SOZ*^*C*^ set (*F*_*SOZ*_ and 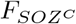 in Figure S1). Then each set of electrodes’ feature distributions were summarized with its quantile statistics over time, resulting in twenty signals: ten quantiles from 10-100 of the SOZ and ten quantiles of the *SOZ*^*C*^ over time. We used a Random Forest (RF) classifier [30]. Specifically, it was a variant known as the Structured Random Forest, or Manifold Random Forests [31, 32]. The manifold RF approach allows one to encode structural assumptions on the dataset, such as correlations in time of the fed in data matrix. The input data matrix for each RF is a multivariate time series that summarizes the quantile statistics of the *SOZ* and *SOZ*^*C*^ over time. As a result, we obtained better results using the manifold RF because it took advantage of local correlations in time, rather than treating all values in the data matrix input as independent as done in a traditional RF model. This approach allowed our classifiers to learn faster with less data, compared to treating all inputs as independent in the traditional RF model. For more information on how manifold RF improves on traditional RF, we refer readers to [31, 32]. For every model, we used the default parameters from scikit-learn and the rerf package [23, 32]: 500 estimators, max depth is None, minimum samples split is 1, max features is auto, feature combinations is 1.5, image height of 20 (20 quantiles total), patch height max of 4, patch height minimum of 1, patch width max of 8, patch width minimum of 1. The output of the decision tree classifier is:

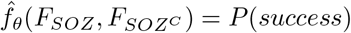

where *θ* are the trained RF parameters and *F*_*SOZ*_ are the heatmap values for the *SOZ* and 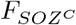 are the heatmap values for the *SOZ*^*C*^ for a specific feature representation F (shown in Supplemental Figure S1). 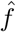 is the function we are trying to estimate, which predicts the probability of successful surgical outcome for a given feature heatmap.

#### Hyperparameters

When looking at iEEG data, clinicians inherently select windows of interest due to external and prior information about when the seizure clinically manifests. Similarly, we consider a window of 10 seconds before seizure onset to the first 5% of the seizure event. This window was chosen apriori to analysis, and we repeated the analyses with slightly varying windows, but the results were consistent. To provide further contrast to the spatiotemporal heatmaps, we considered also a threshold between 0.3 and 0.7 (spaced by 0.1) that would be applied to the heatmap such that values below were set to 0. This is analagous to clinicians being able to look at a iEEG data and hone in only on areas that are extreme relative to the rest. We selected a fixed threshold through nested cross-validation, where thresholds are selected on the train/validation set, and then performance is measured on the held out test set.

#### Structured Heatmap Input

To train a RF classifier on patients with varying number of channels, we structured the spatiotemporal heatmap inputs to summarize the distributions of the feature values in the *SOZ* and *SOZ*^*C*^ sets. We converted these into a vector of 20 quantile values (10 quantiles for *SOZ* and 10 for *SOZ*^*C*^), taken evenly from the 10th to the 100th quantiles. This forms a data matrix per heatmap, which summarizes the *SOZ* and *SOZ*^*C*^ distributions over time, fixed around the seizure onset time.

#### Nested Cross-Validation Feature Evaluation

It is common practice when building a supervised machine learning model to incorporate cross-validation (CV) where the dataset is split into a training and testing set. However, when one has hyperparameters (in addition to the machine learning model parameters) it is crucial to control against over-fitting to the test dataset [33]. Due to our hyperparameter selection of the optimal heatmap thresholds, we used a nested CV scheme, where we split the dataset into a training, validation and a “lock-box” dataset, where the hyperparameters were tuned on an inner CV (70% of the dataset) with the training and validation data, and then performance was evaluated on the lock-box dataset. All features were optimized separately, so that their hyperparameters were optimal for that feature. We computed the discrimination statistic known as the area under the curve (AUC). We repeated the nested-cross validation 10 times, resulting a 10-fold CV. Statistical analysis was done using nested cross-validation, splitting training, validation and testing groups based on subjects, while sampling proportionally the different epilepsy etiologies. We set aside 60% of subjects for training, 10% for validation and then 30% as a held-out test set. We used a heuristic of attempting to sample approximately randomly from clinical complexity 1-4 as defined in the paper. This heuristic attempted to place at least 25% of the subjects in the training, validation and held-out test set for each clinical complexity. Besides this covariate, the training, validation and held-out test set were determined randomly. We found that this performed slightly better for all features compared to random sampling. We performed patient-level CV, ensuring that no patient was in multiple splits of the dataset (i.e. any one patient is only in the train, test, or lock-box dataset).

#### Statistical analysis - Hypothesis Testing

Success probability values were computed from the RF classifiers trained on the spatiotemporal feature representations of the iEEG data (fragility, spectral features, and graph metrics from correlation and coherence graphs) resulting in a distribution of probabilities for each feature.

To compare the RF model performance across the 20 proposed features of fragility, spectral power and graph metrics, we computed Cohen’s D effect size differences between the groups of interest, and then Mann-Whitney U tests for unpaired data, and paired Wilcoxon rank-sum tests for paired data [20]. We corrected for any multiple comparisons using the Holm-Bonferroni step-down method. In some cases, we also present the Likelihood Q-ratio test (LQRT) results to show that the results are robust to statistical test chosen. The LQRT has been shown to be more robust compared to both the Mann-Whitney U-test and t-test in the presence of noise [34, 35]. LQRT utilizes a bootstrapping procedure, so it’s resolution is 0.001 (i.e. if it produces a p-value of 0, then that means it is < 0.001). Since the Mann-Whitney, Wilcoxon and LQRT tests do not rely on a parametric assumption of normality, no explicit parametric assumptions were made. In certain cases, where we report results of a t-test as well, we assumed that the data distribution might be normal, but this was not formally tested.

We compared the success probabilities stratified by different clinical covariates: surgical outcome, Engel class, clinical complexity, handedness, gender, onset age, and surgery age. We then estimated the effect size differences between distributions in the form of a Cohen’s d statistic. Cohen’s d was estimated using a non-parametric bootstrapping on the observed data with 5000 re-samples used to construct a 95% confidence interval [20]. The null hypothesis of our experimental setup was that the success probabilities came from the same population. The alternative hypothesis was that the populations were different (i.e. a feature could distinguish success from failed outcomes, or between different Engel classes).

### 8.9 Feature evaluation using predicted probability of successful surgery

The fragility and all baseline features proposed generated a spatiotemporal heatmap using EEG snapshots of ictal data (outlined in Figure S1). To compare spatiotemporal heatmaps across features, we computed a probability of success (i.e. a confidence score) that is hypothesized to be high for success outcomes and low for failures for “good” features. We expected that fragility would follow a trend of decreasing confidence as CC, Engel score and ILAE score increase. For each clinical covariate group, we measured the effect size difference via bootstrapped sampling, and the statistical p-value between the distributions. We **hypothesized that**: i) fragility would have an effect size difference significantly different from zero when comparing success vs failed outcomes, ii) in addition, this effect size would correlate with meaningful clinical covariates, such as CC and Engel class and iii) both the effect size and p-value would be better than the proposed baseline features.

The higher the probability of success (closer to 1), the more likely the feature indicated a successful surgery, and the lower it was (closer to 0), the more likely the feature indicated a failed surgery. To compute this value, we first partition the heatmap into a *SOZ* and *SOZ*^*C*^ as seen in Supplemental Figure S1. This forms the two sets of signals that represent the spatiotemporal feature values of the *SOZ* set vs the *SOZ*^*C*^ set of electrodes. Then we take windows of interest, where clinicians find most valuable for *SOZ* localization: the period before seizure and right after seizure onset, and performing a nested CV of RF models. The efficacy of each proposed feature is evaluated based on how well the trained RF model is able to predict the surgical outcomes. We tested our hypotheses stated above by computing a probability of success from each feature heatmap for each patient, and estimated the distribution differences of the CS between various clinical covariates.

### 8.10 Spatiotemporal feature heatmap interpretability

To determine how valid the output probability values are, we first computed a calibration curve, which told us how well-calibrated the probability values were. We furthermore compared the calibration curves across neural fragility, spectral features and graph metrics. With relatively good discrimination measured by the AUC and good calibration, we were able to then compare the CS across clinical complexity (CC) scores, Engel scores and ILAE scores. Since epilepsy and reasons for failed surgeries are so complex, these are clinical methods for stratifying patient groups based on observed etiology. We analyzed how the success probability differs across each of these categories. In addition, we looked at how the success probabilities might differ across other clinical variables, such as sex (M vs F), handedness (R vs L), epilepsy onset age (years), and age during surgery (years).

Qualitatively reading off a spatiotemporal heatmap is highly interpretable as one can match raw iEEG segments to certain time periods in the heatmap. For every patient, the time-varying quantile signals (from 10th to the 100th) of the *SOZ* and *SOZ*^*C*^ were computed for every iEEG heatmap, as specified in Methods - Experimental design. We used permutations feature importance sampling to obtain a relative weighting of each signal over time and how important it was in allowing the RF to correctly determine surgical outcome. This was visualized as a heatmap showing the mean and std importance scores of the *SOZ* and *SOZ*^*C*^ statistics, as shown in Figure S7.

In addition, we claim that human interpretability of the spatiotemporal heatmap relies on contrast between the *SOZ* and *SOZ*^*C*^ regions in successful outcomes. Using the results of the feature importance permutation test, we computed for every heatmap, an interpretability ratio, which was defined as:

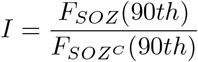

where I is the interpretability ratio for a specific subject’s feature heatmap F. *F*_*SOZ*_(90*th*) is the 90th quantile and up feature values of the *SOZ*, and 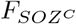 (90*th*) is for the *SOZ*^*C*^. This ratio is then stratified according to surgical outcome, which we expect higher ratios for successful outcomes and lower ratios for failed outcomes. We quantified this difference using Cohen’s D effect size and Mann Whitney U-test Pvalues (see Methods - Statistical analysis - Hypothesis Testing).

### 8.11 Fragility heatmaps are more interpretable than all other EEG feature maps

We can parse the RF model to determine important aspects of the spatiotemporal heatmap that are important in predicting surgical outcome. To do so, we perform permutations on the heatmap inputs to the RF models to measure their relative importances over space and time. We observed that the AUC metric was affected primarily by a combination of the highest (90th quantiles and up) neural fragility values of both the *SOZ* and *SOZ*^*C*^ contacts (Figure S7). The *SOZ*^*C*^ neural fragility as much as 10 seconds before seizure onset impacted the AUC as did neural fragility of the *SOZ* right at seizure onset. For both the *SOZ* and the *SOZ*^*C*^ fragility distributions, the 80th quantile and below did not contribute to the predictive power. In different patients across different clinical centers, the practice of marking seizure onset times can vary due to varying methodologies [5]. As a result, in the *SOZ*^*C*^, there was some variation in terms of which time point in the heatmap mattered most.

Although strong predictive capabilities of an EEG marker of the *SOZ* are promising, it is also important that the marker be presented in an interpretable manner to clinicians. We next show how fragility heatmaps are the most interpretable over all baseline features. In Figure S9a, there are two heatmaps computed for the same seizure event in Patient_01: one is a beta band map (left) and one is a neural fragility map (right). Both maps are normalized across channels and both are computed with similar sliding windows. However, less contacts “stand-out” as pathological in the beta-band heatmap before seizure onset, with the majority of the map being different shades of blue. In contrast, in the fragility heatmap, one contact (**ATT1** from Figure 4) is fragile the entire duration before seizure onset (solid white line), and then a few more contacts become fragile after the electrographic onset of the seizure. These fragile areas that “pop-out” in the heat-map as red-hot areas occur in the clinically annotated SOZ and this patient had a successful surgery.

To quantify interpretability, we compute an interpretability ratio: the ratio of the feature values in the 90th quantile between the *SOZ* and the *SOZ*^*C*^ over the section of data used by the feature’s RF model. This measures the contrast that one sees between the extreme values in the *SOZ* versus the extreme values they see in the *SOZ*^*C*^. The larger the ratio, then the more contrast the map will have. In Figure S9b, we show that the effect size difference between successful and failed outcome of this interpretability ratio is largest in neural fragility when compared to all baseline features. It is well-known that RF models are scale-invariant [30], so it is plausible that there are portions of heatmaps that distinguish one channel from another that can be parsed out via a decision tree and not the naked eye, which leads to high AUC in the beta band representation in Figure 5. For example a decision tree can discriminate between 0.3000 and 0.2995, which on a normalized color-scale is difficult to parse by visual inspection. Neural fragility on the other hand, shows marked differences between the clinically annotated *SOZ* (red electrodes on the y-axis) and the actual fragility values even before seizure onset.

## 9 Extended Data Figures

**Figure S1:**
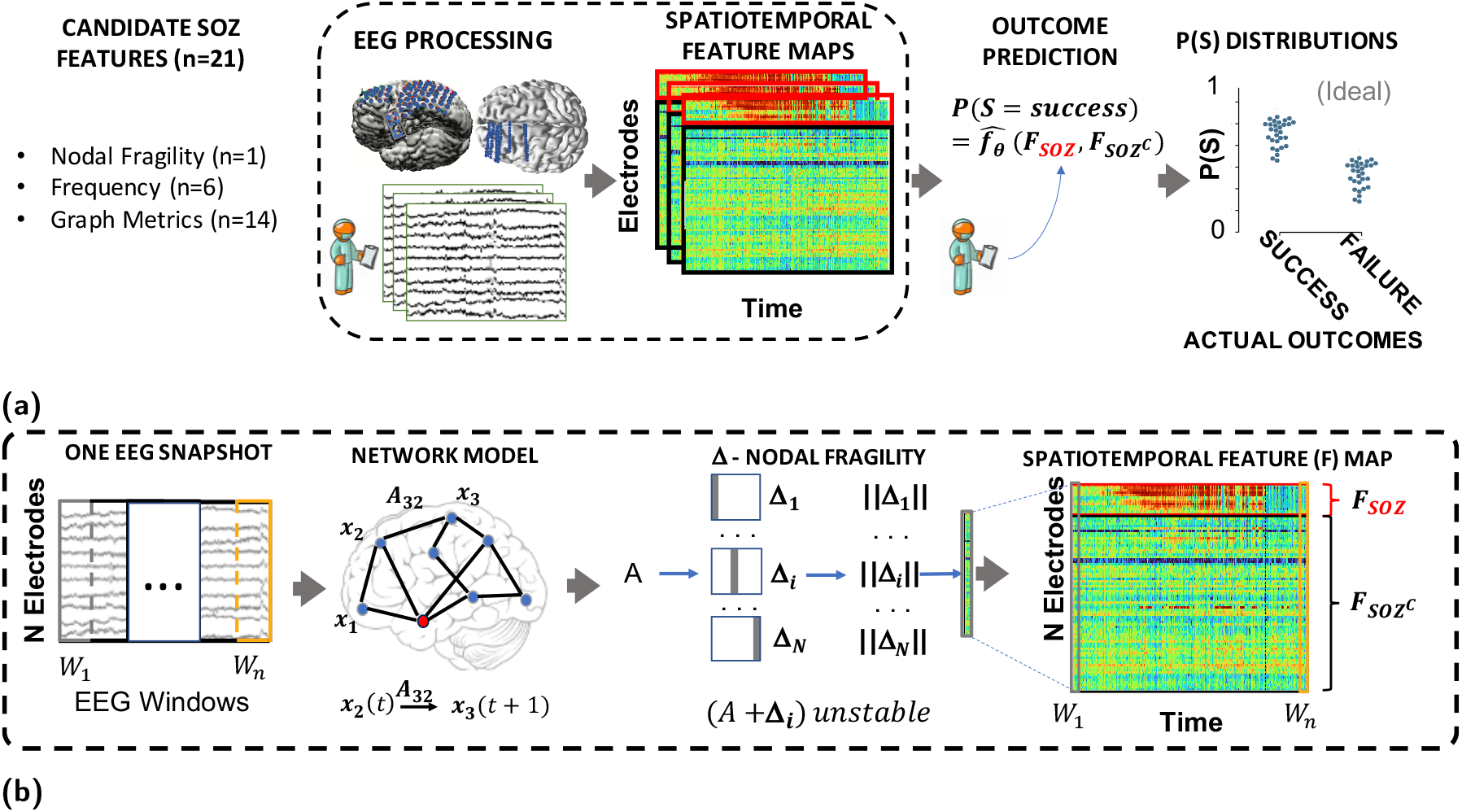
Computational experiment setup for all candidate *SOZ* features and statistical analysis. - **(a)** Any candidate feature that can produce a spatiotemporal heatmap was computed from EEG data and then partitioned by the clinically annotated *SOZ* set and the complement, *SOZ*^*C*^ (i.e. non-*SOZ* electrodes) to compute a confidence statistic measuring the feature’s belief of the clinician’s hypothesis. Here *F*_*SOZ*_ and 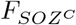 were the feature values within their respective sets. *f*_*θ*_ is the function depending on the Random Forest model parameters, *θ* that maps the statistics of the *F*_*SOZ*_ and 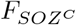 to a confidence statistic. An ideal feature would have high and low confidence for success and failed outcomes respectively. Each point on the final CS distribution comparisons represent one patient. **(b)** A more detailed schematic of how our proposed fragility and baseline features were computed from EEG data for a single snapshot of EEG data. See fragility methods section for description of x, *A* and Δ.

**Figure S2:**
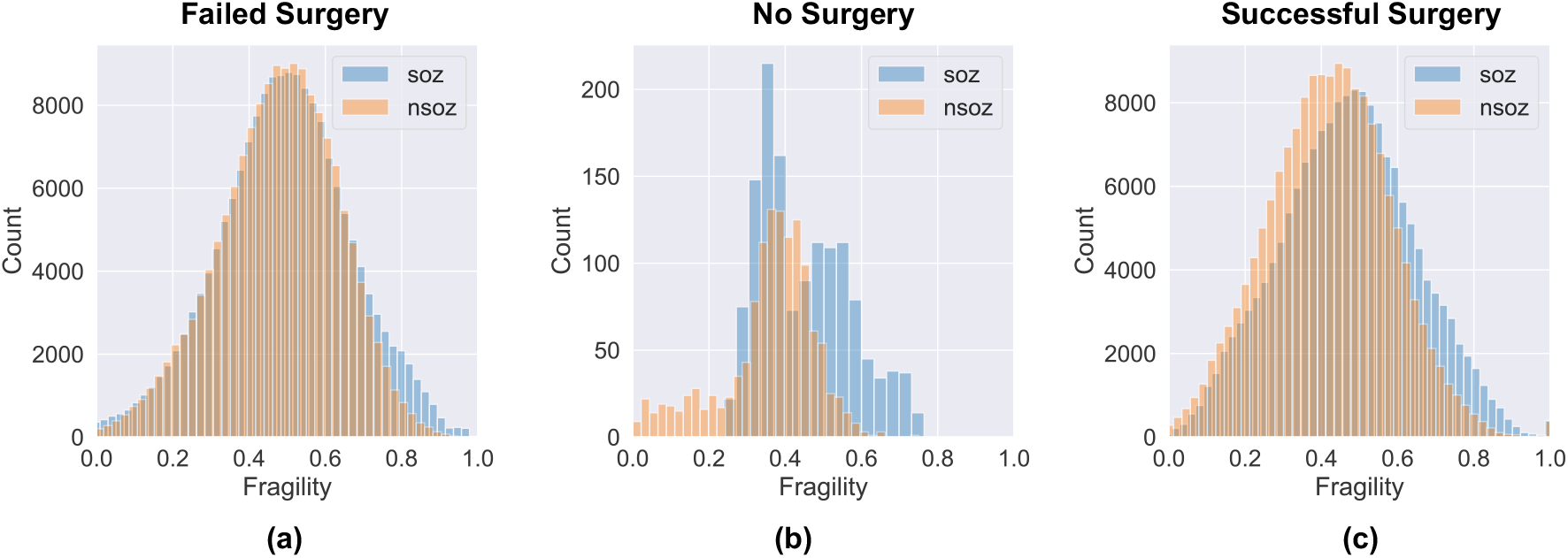
Pooled fragility distribution analysis for all patients. - failed **(a)**, no surgery **(b)** and successful surgery **(c)** datasets. Each *SOZ* (soz in blue bars) and *SOZ*^*C*^ (‘nsoz’ in orange bars) distribution per patient was bootstrap sampled (see Methods for more information on sampling) and then compared using the one-sided Mann-Whitney U test. The corresponding test yielded a statistic of 2776334 (PValue = 0.355) for the failed patient outcomes and a statistic of 36836739 (PValue = 3.326e-70) for the successful patient outcomes. The patients without resection were not included in the analysis comparing to outcome, but these patients can present as interesting case studies where the SOZ was hypothetically localizable, but perhaps was too close to eloquent areas.

**Figure S3:**
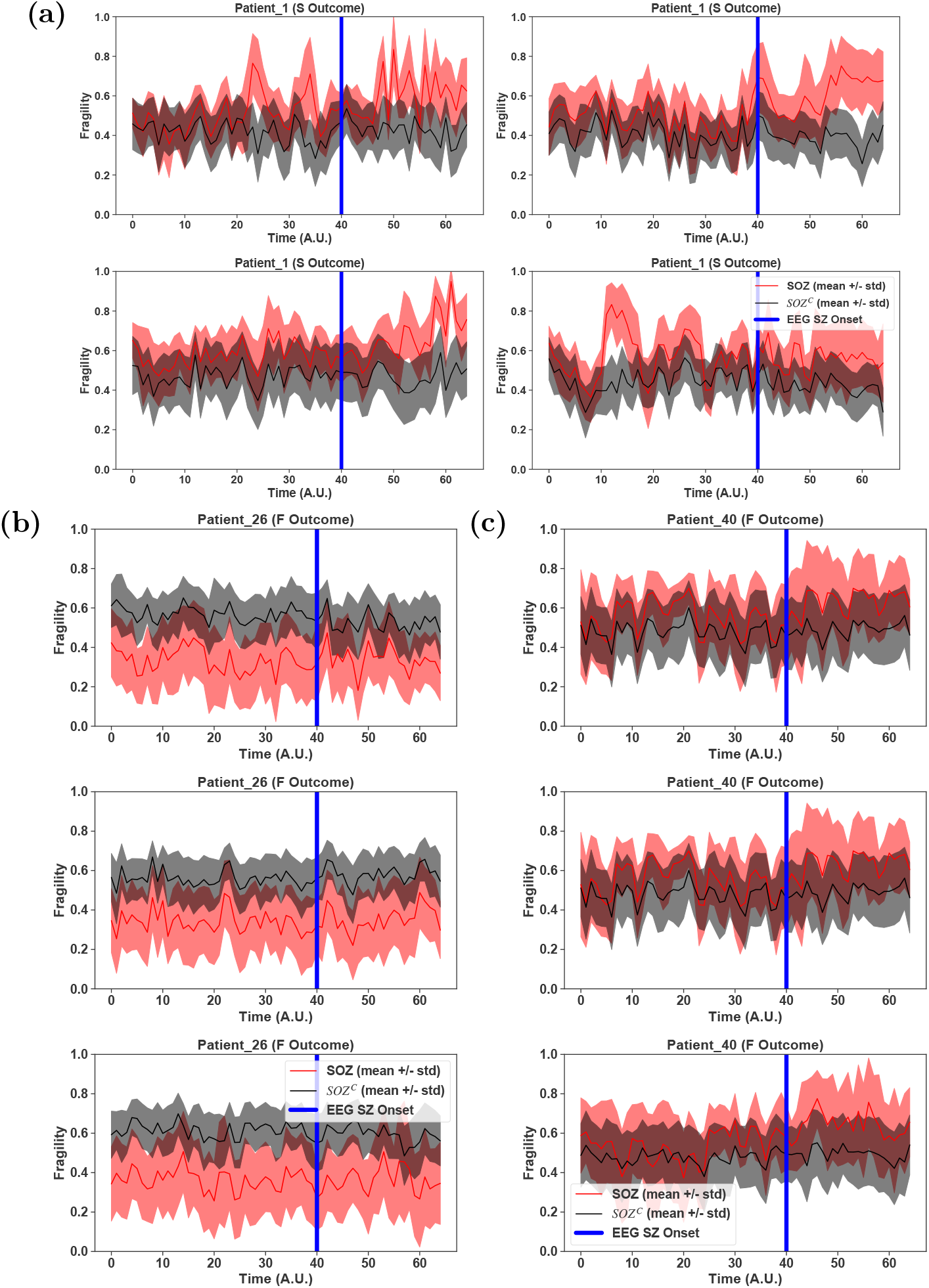
Patient-specific *SOZ* vs *SOZ*^*C*^ neural fragility near seizure onset. - Red *SOZ* vs black *SOZ*^*C*^ signals for patients presented in Figure 4: Patient_01 **(a)**, Patient_26 **(b)**, Patient_40 **(c)**. For each patient, the ictal snapshots available are visualized around seizure onset with 5 seconds before onset until the first 20% of the seizure. Not necessarily all electrodes in the clinically annotated *SOZ* are part of the EZ when the patient had a successful outcome. Therefore, if neural fragility had value in contrasting true EZ electrodes from non-EZ electrodes, then any extra electrodes clinically annotated in the *SOZ* should have relative lower fragility. The lines represent mean +/−sem.

**Figure S4:**
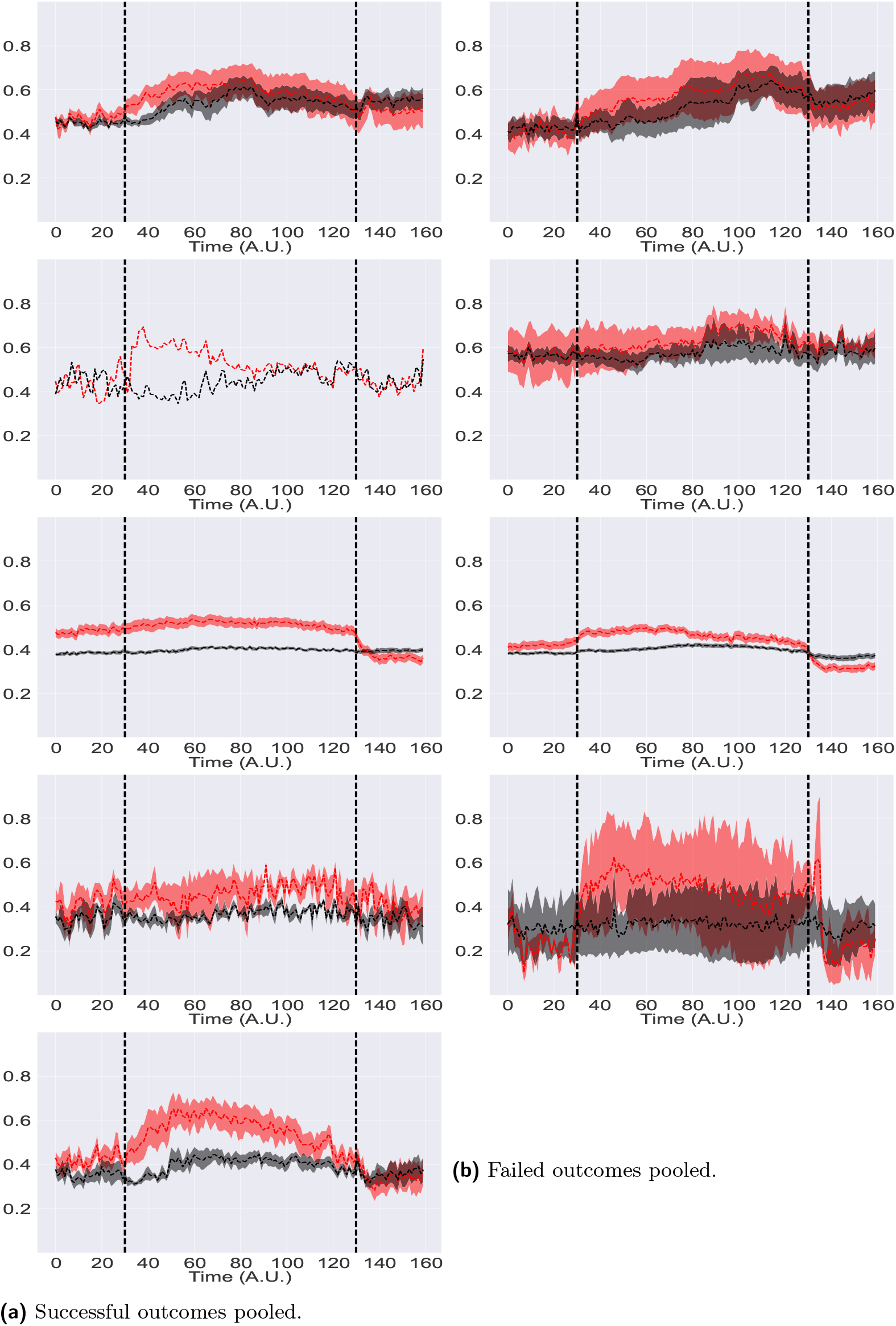
Pooled-patient per clinical center *SOZ* vs *SOZ*^*C*^ neural fragility. - Red *SOZ* vs black *SOZ*^*C*^ fragility signals for pooled patients within each of the five centers with successful **(a)** and failed outcomes **(b)** for NIH (n=14), JHH (n=4), CC (n=61), UMH (n=5), and UMMC (n=7) (top to bottom respectively). Note UMMC only had successful outcomes, so there was no curve for the failures. Seizure periods were resampled and normalized to 100 samples for averaging and viewing purposes. In JHH and UMH, there were only one and two patients in successful outcomes respectively. The lines represent mean +/−sem.

**Figure S5:**
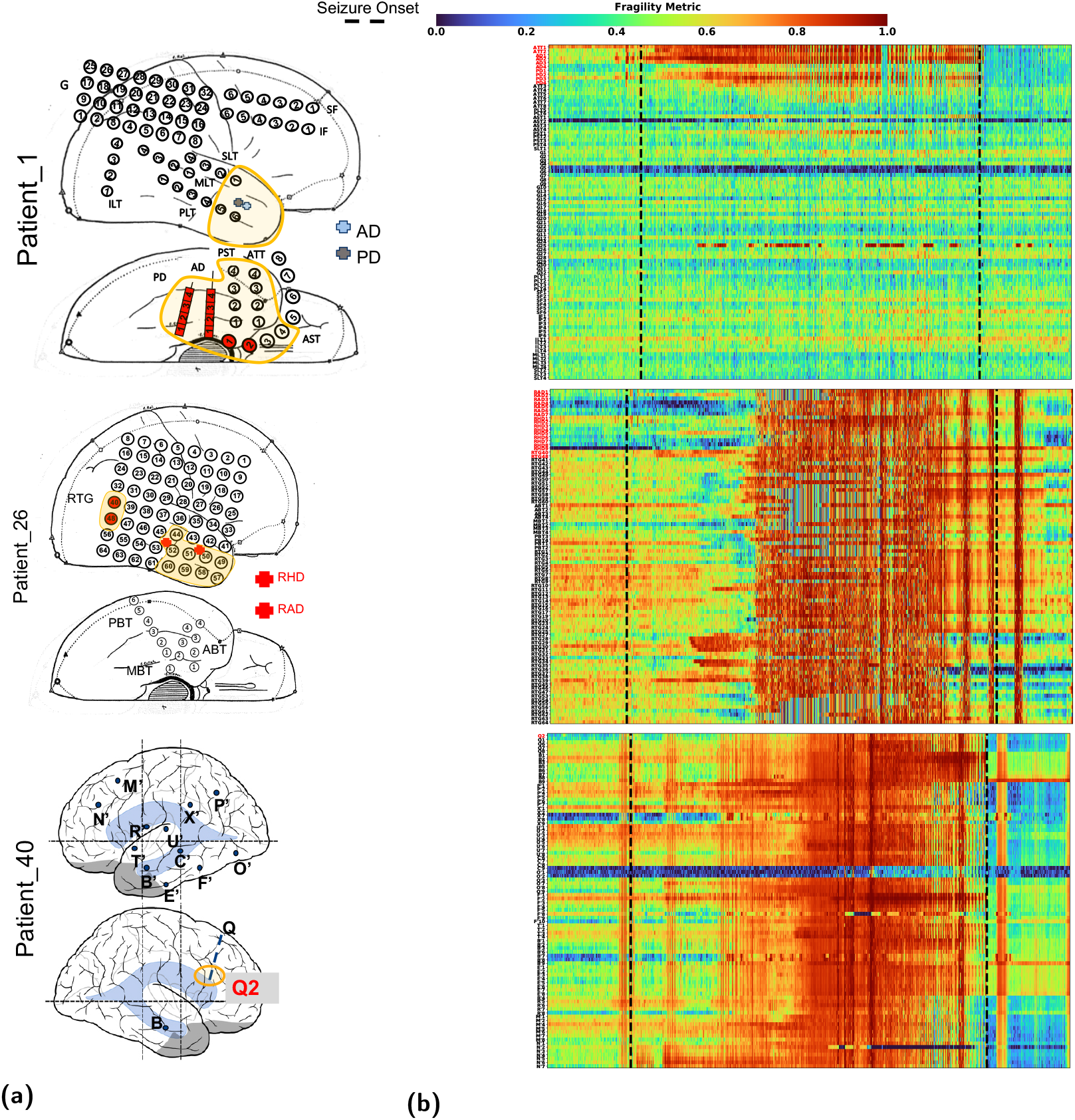
Entire fragility heatmap of seizures in successful and failed surgical outcomes. - Fragility heatmaps with electrodes on y-axis and time on x-axis with the dashed white-lines denoting seizure onset and offset. Shows a period of 30 seconds before seizure onset and 30 seconds after seizure offset. **(a)** Shows clinically annotated maps of the implanted ECoG/SEEG electrodes with red denoting SOZ contacts. **(b)** shows spatiotemporal fragility heatmaps for example of successful outcome (Patient_01), and failed outcome (Patient_26 and Patient_40). The color scale represents the amplitude of the normalized fragility metric, with closer to 1 denoting fragile regions and closer to 0 denoting relatively stable regions. The contacts in red and orange are part of the SOZ and RZ, respectively as defined in Methods section. Note that the red contacts are also part of the RZ. Within the seizures, estimating the linear systems are not as stable, which can be seen by fragility “everywhere” in the map. Visualized with Turbo continuous colormap. Best seen if viewed in color.

**Figure S6:**
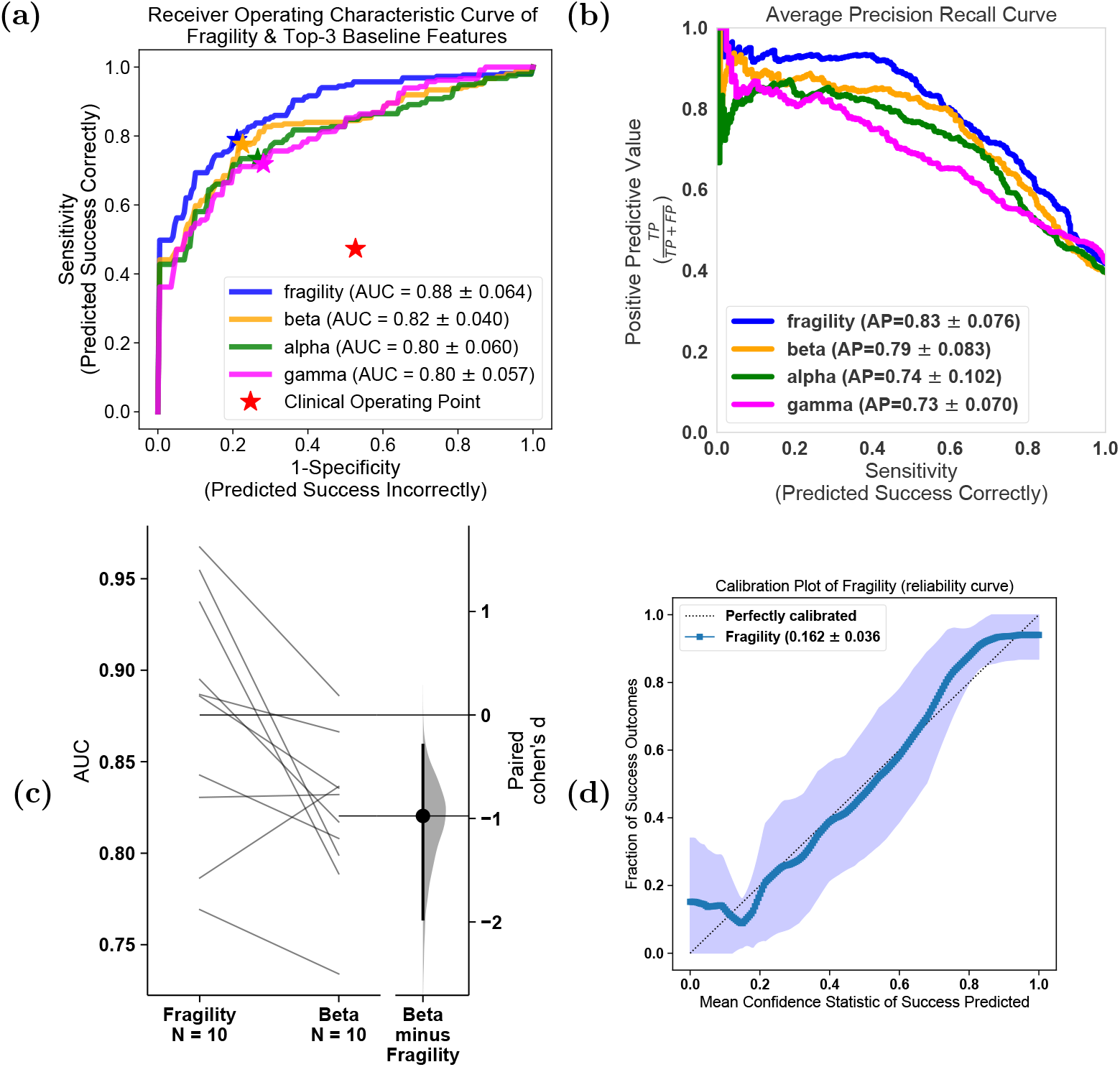
Comparison of classification models using different features. - **(a)** The ROC curve over 10 folds of cross-validation of the held-out test set obtained by applying a Random Forest model onto the spatiotemporal heatmaps to predict surgical outcome (see Methods). Fragility and the top-3 baseline features in terms of AUC are visualized. The shaded area represents the standard deviation of the curve obtained by linear interpolation for visualization purposes. The AUC of fragility obtained a 0.88 +/−0.064 over the 10 standard deviation with a relative improvement of 7.2% improvement in AUC compared to the next best feature representation (i.e. the beta frequency band). At the Youden point (stars), neural fragility obtains a balanced accuracy score of 0.76 +/−0.06, and an improvement of 0.32 in TPR and 0.32 in FPR compared to the clinical operating point (red star). **(b)** The average PR curve showing that fragility is better then the top 3 features by at least an average precision of 0.04. **(c)** A paired estimation plot showing how the same test set of patients differed in AUC depending on whether it was using the fragility, or beta feature heatmap representation. The paired Cohen’s D effect size was computed at −0.975 (−1.97 to −0.29; 95% CI). The p-values associated with the difference between Neural Fragility and the Beta frequency band were 0.0204, 0.0273, and 0.0225 using the one-sided Wilcoxon rank-sum test, permutation test, and the paired student t-test respectively. **(d)** Calibration curve showing the fraction of actual successful surgical outcomes on the y-axis vs the average CS output on the x-axis. The curve measures how calibrated the predicted success probability values are to the true risk stratification of the patient population. The closer a curve is to the *y* = *x* line, then the more calibrated a model is. It is quantified by the Brier-loss (closer to 0 is better), which is shown in the legend, and is significantly lower then the next best feature (an improvement of 15%). The shaded region represents 95% confidence interval of two standard deviations.

**Figure S7:**
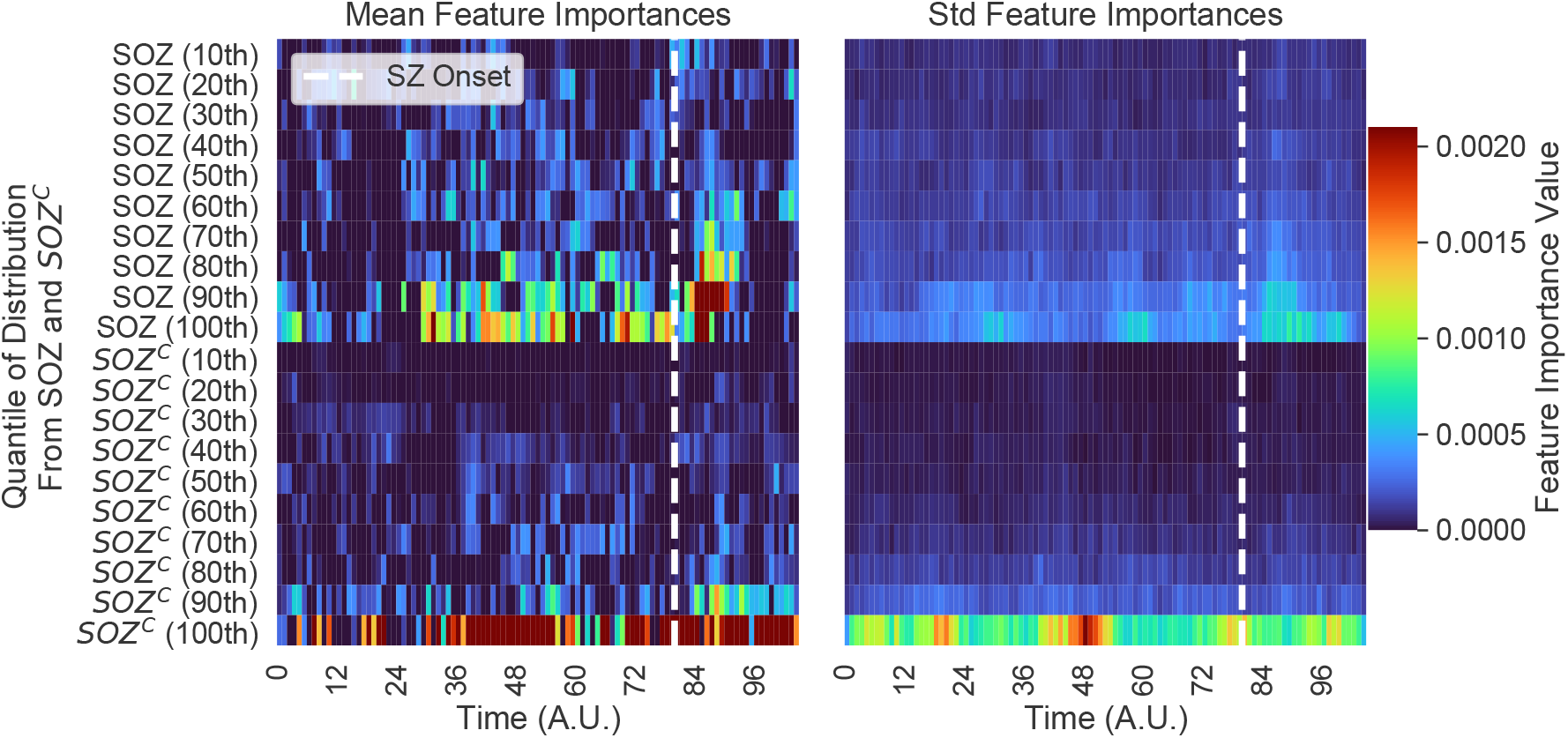
Estimated feature importance (mean and stdev) of the associated fragility heatmap used estimated using permutation. - The metric of interest was the concordance statistic (i.e. AUC) of the ROC curve. The original feature map is transformed into a 20-dimensional set of time-varying statistics of its *SOZ* and *SOZ*^*C*^ electrodes describing the quantiles of the spatiotemporal heatmap (10% - 100% quantiles). This time-varying summary allows these heatmaps to be pooled together across subjects when training a Random Forest classifier as described in Methods.

**Figure S8:**
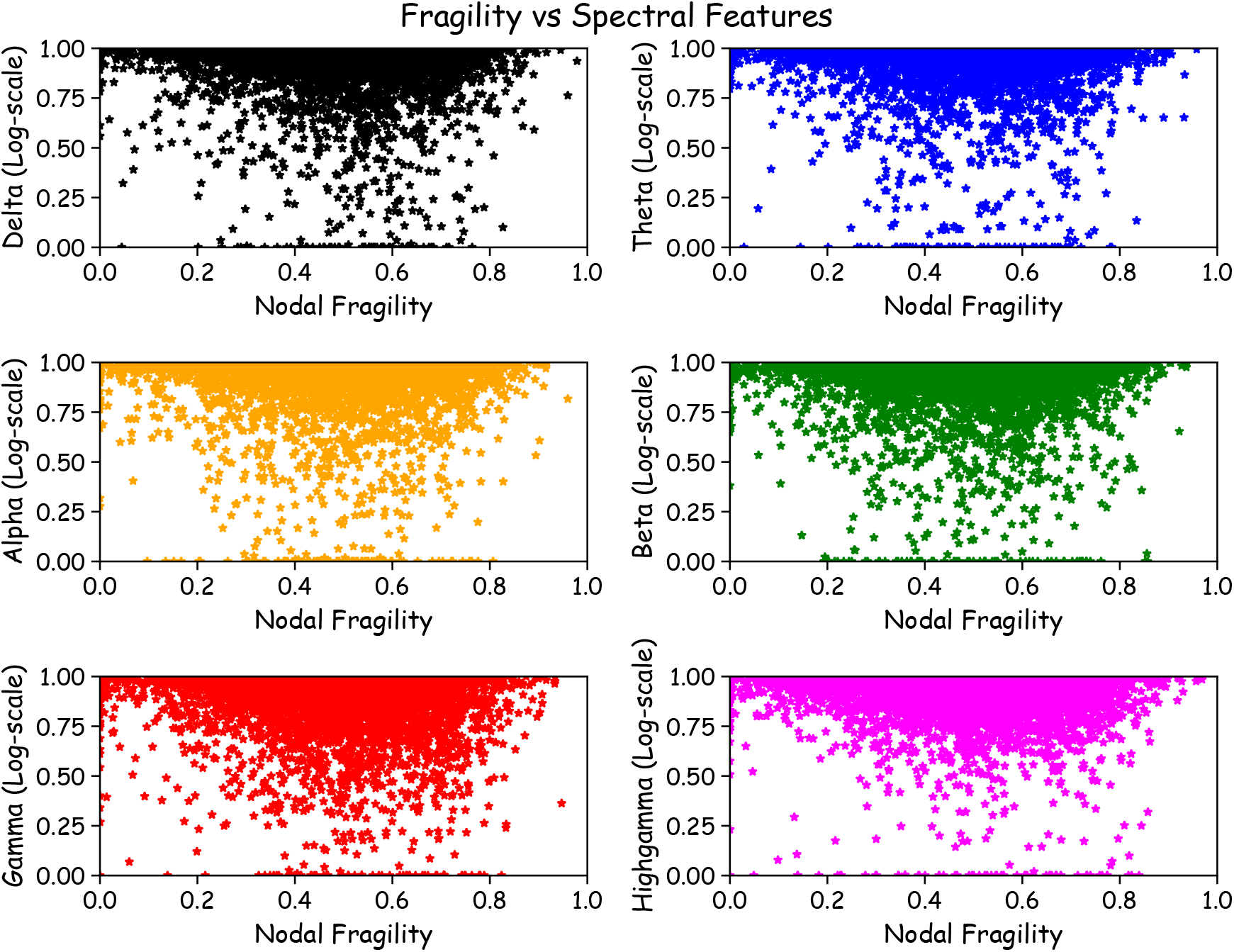
Neural fragility vs frequency power values. - Fragility versus frequency power in the delta, theta, alpha, beta, gamma and highgamma band for Patient_01, Patient_26, and Patient_40. For band definitions, refer to Methods - Baseline features - spectral features. Every point represents the spectral power and neural fragility value from a randomly chosen window and electrode from one of the patients. No significant correlation is seen or computed from the data. Each spectral feature and fragility are normalized as described in Methods.

**Figure S9:**
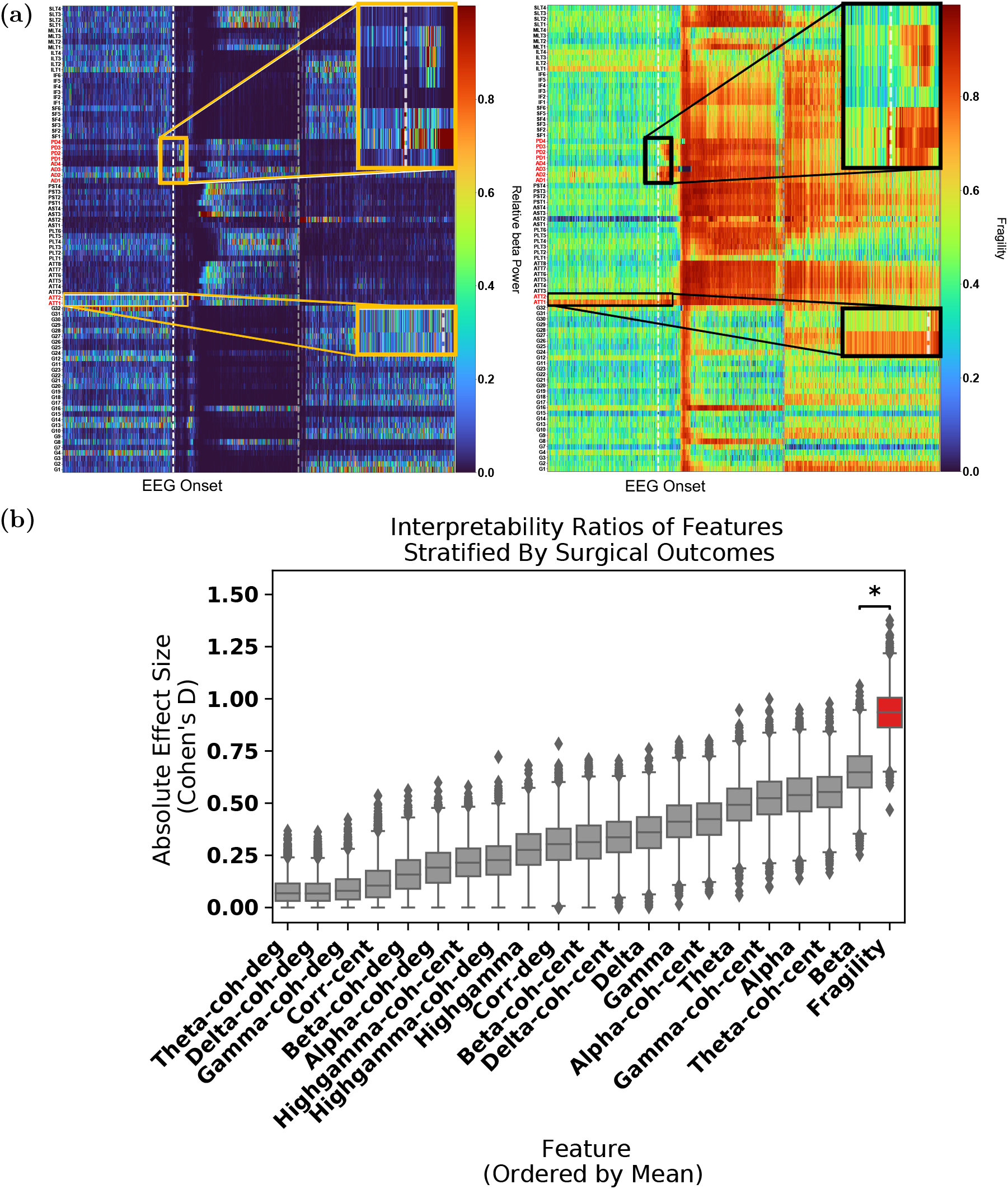
Interpretability ratio of feature heatmaps. **(a)** Two heatmap examples of a seizure snapshot of Patient_01 (NIH treated, ECoG, CC1, Engel I, ILAE 1) with the beta frequency band (left) and the neural fragility heatmap (right). Both colormaps show the relative feature value normalized across channels over time. The black line denotes electrographic seizure onset. **(b)** A box plot of the interpretability ratio that is defined in Results Section 8.11 computed for every feature. The y-axis shows an effect size difference between the interpretability ratios of success and failed outcomes. The interpretability ratio for each patient’s heatmap is defined as the ratio between the feature values in the two electrode sets 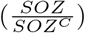. Neural fragility is significantly greater then the beta band (alpha level=0.05).

**Figure S10:**
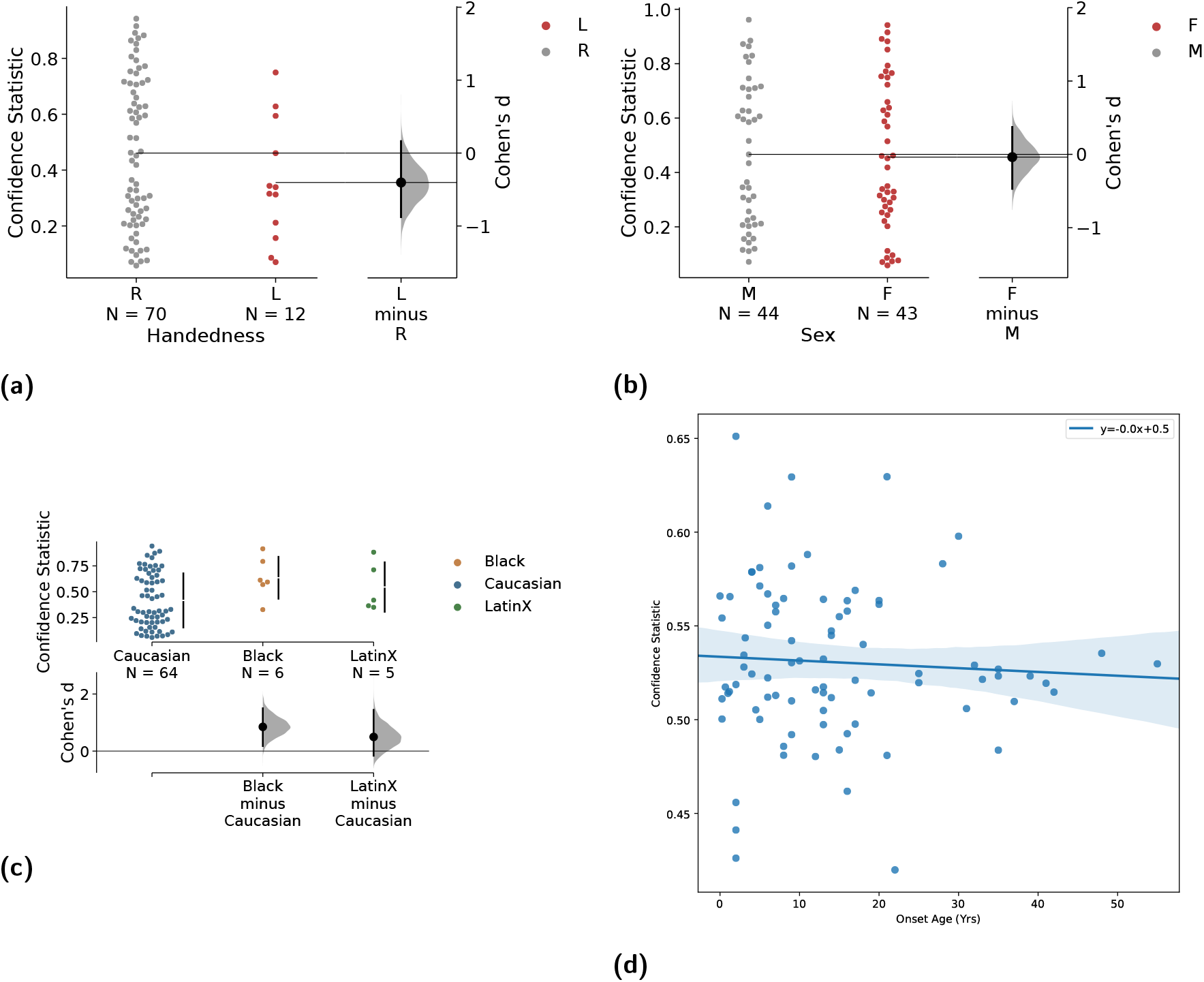
Neural fragility correlation against non-epileptic clinical covariates. - Fragility success probabilities (denoted as “Confidence Statistic” in y-axes) split by clinical factors, such as handedness **(a)**, gender **(b)**, ethnicity **(c)** and age at surgery **(d)**. Not all patients had data for each of these categories, so the subset of available data was used. Note the sample sizes vary across different groups shown. For ethnicity, we also had 1 Asian subject, but left it out because the permutation effect size estimation procedure does not work for 1 sample. Effect sizes were estimated using the permutation test and Mann Whitney U test described in Methods. The corresponding effect sizes and p-values were (0.1/0.99) for handedness, and (0.12/0.7) for gender. The pvalue was computed using the one-sided Mann-Whitney U test. The slope was negligibly close to 0 for surgery age linear fit. There was no relatively significant trend in the data related to ethnicity. The significant Cohen’s D effect size difference is primarily due to the low sample sizes in non-Caucasian ethnicities. The error bars represent 95% confidence interval specified by 2 standard deviations.

